# Comparison of tri-exponential decay vs. bi-exponential decay and full fitting vs. segmented fitting for modeling liver intravoxel incoherent motion diffusion MRI

**DOI:** 10.1101/429977

**Authors:** Olivier Chevallier, Nan Zhou, Jean-Pierre Cercueil, Jian He, Romaric Loffroy, Yì Xiáng J. Wáng

## Abstract

**Purpose:** To determine whether bi- or tri-exponential models, and full or segmented fittings, better fit IVIM imaging signal of healthy livers.

**Materials and methods:** Diffusion-weighted images were acquired with a 3-T scanner using respiratory-triggered echo-planar sequence and 16 *b*-values (0∼800 s/mm^2^). Eighteen healthy volunteers had liver scanned twice in the same session, and then once again in another session. Region of interest (ROI)-based measurements were processed with bi-exponential model full fitting and segmented fitting (threshold *b*-value = 80 s/mm^2^), as well as tri-exponential model full fitting and segmented fitting (threshold *b*-value = 200 s/mm^2^).

**Results:** With all scans’ signal averaged, bi-exponential model full fitting showed D_slow_=1.14, D_fast_=193.6×10^-3^ mm^2^/s, and PF=16.9%, and segmented fitting showed D_slow_=1.03, D_fast_=56.7×10^-3^ mm^2^/s, and PF=21.3%. IVIM parameters derived from tri-exponential model were similar for full fitting and segmented fitting, with a slow (D’_slow_=0.98×10^-3^ mm^2^/s; F’_slow_=76.4 or 76.6%), a fast (D’_fast_=15.1 or 15.4×10^-3^ mm^2^/s; F’_fast_=11.8 or 11.7%) and a very fast (D’_Vfast_=445.0 or 448.8×10^-3^ mm^2^/s; F’_Vfast_=11.8 or 11.7 %) diffusion compartments. Tri-exponential model provided an overall better fit than bi-exponential model. For bi-exponential model, full fitting provided better fit at very low and low *b*-values compared with segmented fitting with the later tended to underestimate D_fast_, however, segmented method demonstrated lower error in signal prediction for high *b*-values. Compared with full fitting, tri-exponential segmented fitting offered better scan-rescan reproducibility.

**Conclusion:** For healthy liver, tri-exponential modelling is preferred than bi-exponential modelling. For bi-exponential model, segmented fitting underestimates D_fast_, but offers more accurate estimation of D_slow_.

## Introduction

Intravoxel incoherent motion (IVIM) reflects the random microscopic motion that occurs in voxels on magnetic resonance (MR) images of water molecules (either intra-cellular or extracellular) and the microcirculation of blood. In 1988, Le Bihan *et al*. proved that it was possible to identify two different compartments of diffusion in the brain (1). According to IVIM theory, the fast component of diffusion (represented by D_fast_, or *D*^*^) is related to micro-perfusion, whereas the slow component (represented by D_slow_, or *D*) is linked to molecular diffusion. The signal decay of IVIM diffusion MRI is therefore described using a bi-exponential model [1]. In recent years, IVIM diffusion MRI has been tested in many studies, and promises potential important clinical values. However, bi-exponential modelling of liver IVIM signal is known to be challenging, with an unsatisfactory reproducibility especially for D_fast_ estimation, owing to the usually limited *b*-value sampling and low signal-to-noise ratio (SNR) with diffusion weighted (DW) images (2, 3). The physiological motion, particularly those associated with respiration, complicates data acquisition during the long scan duration with multiple *b*-values (3, 4). For bi-exponential IVIM model, a segmented approach, that is to calculate D_slow_ firstly by linear regression, then PF by extrapolation, then D_fast_ by nonlinear, has frequently been used in the literature since simultaneous fitting of all diffusion parameters usually gives less stable results (5). However, theoretically it is possible that full fitting results may more resemble physiological value of IVIM parameters.

Liver is histologically unique in many aspects, which include the presence of several vessel types (arteries/arterioles, portal veins/venules, hepatic veins/venules), sinusoid capillaries, bile ducts, a rich lymphatic system, and a functionally important intermediate area between the sinusoids and hepatocytes, called the “space of Disse”. Flowing or moving spins are present in these compartments, which are directly or indirectly connected together. With MR signal of all study participants averaged together and nonlinear least-squares full fitting method, Cercueil *et al*. demonstrated that a third very fast diffusion compartment may exist, though the precise origin of the third compartment cannot be precisely defined; and the tri-exponential model provided the best fit for IVIM signal decay in the liver over the 0–800 s/mm^2^ range (6). More recently, Wurnig *et al*. investigated an extensive DW-imaging protocol including 68 *b*-values and computed apparent diffusion coefficient ‘spectra’, and demonstrated the presence of a third component of diffusion in liver and kidney (7). However, till now tri-exponential decay model has not been studied at individual subject level with clinically applicable examination set-up. With 50 IVIM diffusion MR scans from 18 healthy volunteers, this study analyzes the fitting accuracy of bi-exponential vs. tri-exponential models, and full fitting vs. segmented fitting methods. Fitting residual error of predicted MR signal vs. measured MR signal with the four fitting approaches, as well as scan-rescan repeatability/reproducibility at individual subjects’ level, are compared. An analysis of the error distribution in model signal prediction can offer insight into why tri-exponential model offers better fit than bi-exponential model for liver diffusion signal decay. The IVIM parameter physiological values associated with each fitting approach are also presented in this paper.

## Methods

This study was conducted with the approval of the institutional ethics committee and informed consent was obtained. MRI scans on 18 healthy volunteers (mean age: 25.7 years; range: 24–27 years, males = 5, females = 13) were performed between April and May 7, 2017. All volunteers were scanned twice during the same session (scans 1.1 and scans 1.2), and additionally once again in another session (scan 2) with an interval of 5–21 days (mean 13 days). DW imaging was acquired with a 3T magnet and a 32 channels dStream Torso coil (Ingenia, Philips Healthcare, Best, The Netherlands). The IVIM diffusion imaging was based on a single-shot spin-echo-type echo-planar imaging sequence, with 16 *b*-values of 0, 3, 10, 25, 30, 40, 45, 50, 80, 200, 300, 400, 500, 600, 700 and 800 s/mm^2^, NSA of 2 for *b* = 700 s/mm^2^ and *b* = 800 s/mm^2^, and NSA = 1 for other *b*-values. Spectral presaturation with inversion recovery technique was used for fat suppression. Respiratory triggering was performed using an air-filled pressure sensor fixed on the upper abdomen, resulting in an average TR of 2149 ms. Other parameters included TE = 55ms, slice thickness = 6mm, matrix = 100×116, field-of-view (FOV) = 360×300 mm, EPI factor = 29, a sensitivity-encoding (SENSE) factor = 4, number-of-slices = 26. The study subjects were trained so that they maintained a gentle regular breathing during image acquisition. The average IVIM scan duration was 6 min.

As described previously (4), for the acquired image data, a manual procedure was taken to ‘clean the image data’ for each examination in order to remove image series contaminated by evidential motion and other artifacts [Fig 1]. Four subjects each had one scan with insufficient image quality (4), therefore finally 50 scans were included for models comparison, and 17 scan pairs and 14 scan pairs for repeatability and reproducibility assessment respectively [Fig 1]. Among these 50 scans, the slice number utilized for analysis was on average 7.6 (median: 7, range: 3–12 slices). Curve-fitting algorithms were implemented in a custom program developed on MATLAB (Mathworks, Natick, MA, USA). Regions-of-interest (ROIs) were then placed to cover a large portion of right liver parenchyma while avoiding large vessels on *b* = 0 s/mm^2^ images of the selected image series, then copied and pasted on each corresponding image of each *b*-values. The mean signal intensity of each ROIs was weighted by the number of pixels included in each ROI, then the average of the weighted mean signal intensity of each ROI was calculated to obtain the average signal value of the liver. The signal value at each *b*-value was normalized by attributing a value of 100 at *b*=0 s/mm^2^ (S_norm_ = (SI / SI_0_) × 100, where S_norm_ is the normalized signal, SI = signal at a given *b*-value, and SI_0_=signal at *b*=0 s/mm^2^).

**Figure 1.**
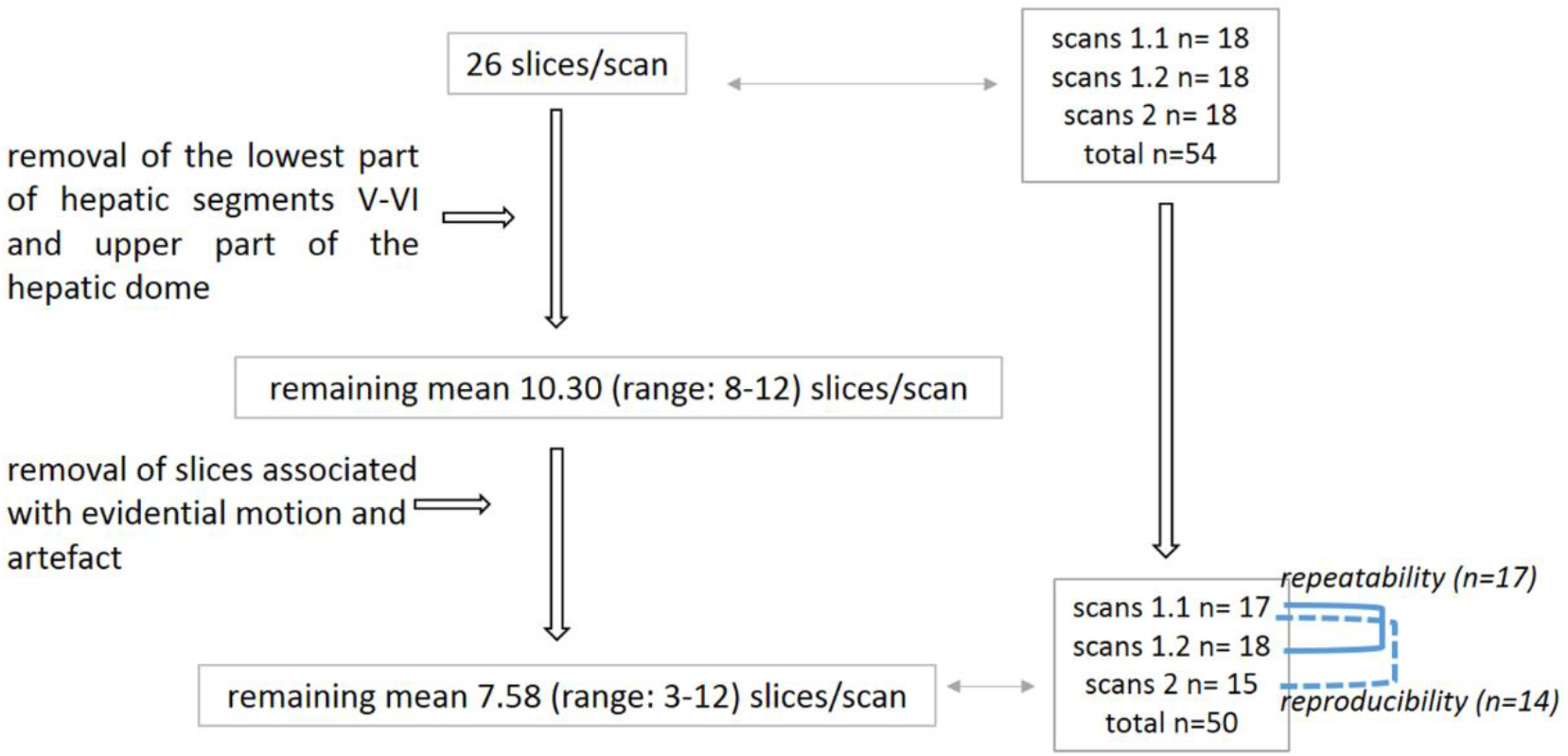
Image-data cleaning process flow diagram and subjects/scans number included for analysis.

For bi-compartmental model, the signal attenuation was modeled according to Eq 1 (8): 

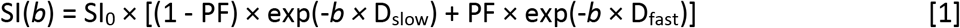

where SI(*b*) and SI_0_ denote the signal intensity acquired with the *b*-factor value of *b* and *b*=0 s/mm^2^, respectively (8). The perfusion fraction (PF) represents the fraction of the pseudo-diffusion compartment related to microcirculation, Dslow is the diffusion coefficient representing the slow (pure) molecular diffusion, and D_fast_ is the pseudo-diffusion coefficient representing the incoherent microcirculation within the voxel (perfusion-related diffusion).

For tri-compartmental model, the signal decay was modeled according to following Eq 2 (6):

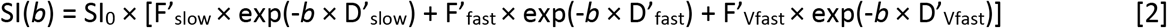

Where F’_slow_, F’_fast_ and F’_Vfast_ are the fractions of each compartments (F’_slow_ + F’_fast_ + F’_Vfast_= 1), meaning that F’_slow_ and (F’_fast +_ F’_Vfast_) are respectively similar to the fractions of the true diffusion compartment (1 – PF) and to PF of the bi-exponential model. D’_slow_ represents the diffusion coefficient, thus similar to D_slow_, D’_fast_ and D’Vfast represent the fast and very perfusion-related pseudo-diffusion coefficients (6). Additionally, F’_Vfast_= 1 - F’_slow_ - F’_fast_ is assumed so to simplify the equation with 5 rather than 6 parameters.

With full fitting method, all the parameters (D_slow_, D_fast_, PF, and D’_slow_, D’_fast_, D’_Vfast_, F’_slow_, F’_fast_) were estimated by a single least-squares nonlinear regression. For bi-exponential model segmented fitting, the threshold *b-T* value was chosen to be *b*=80 s/mm^2^ (4, 9). For tri-exponential segmented fitting, the estimation of D’_slow_ was obtained by a least-squares linear fit of the logarithmized image intensity with *b*-values ≥ 200 s/mm^2^ to a linear equation (4) (supplement document 1). The obtained D’_slow_ was then substituted into Eq. 3 and a nonlinear least-squares fit against all *b* values estimated D’_fast_, D’_Vfast_, F’_slow_, F’_fast_. Trust-Region based algorithm was used for all nonlinear regressions, allowing boundary constraints, which is preferable in order to get plausible values when SNR is low (10). The starting points and boundary constraints for bi- and tri-exponential model are shown in Table 1 (supplement document 2). The coefficients of determination R^2^ and adjusted-R^2^ were calculated for each DW image datasets in order to quantify the goodness of the fits. Adjusted-R^2^ is a better indicator when comparing nested models, since its calculation takes into account the number of parameters (11) (supplement document 3). Using ROI-analysis, a R^2^ value higher than 0.95 generally indicates good fit (12, 13).

**Table 1.**
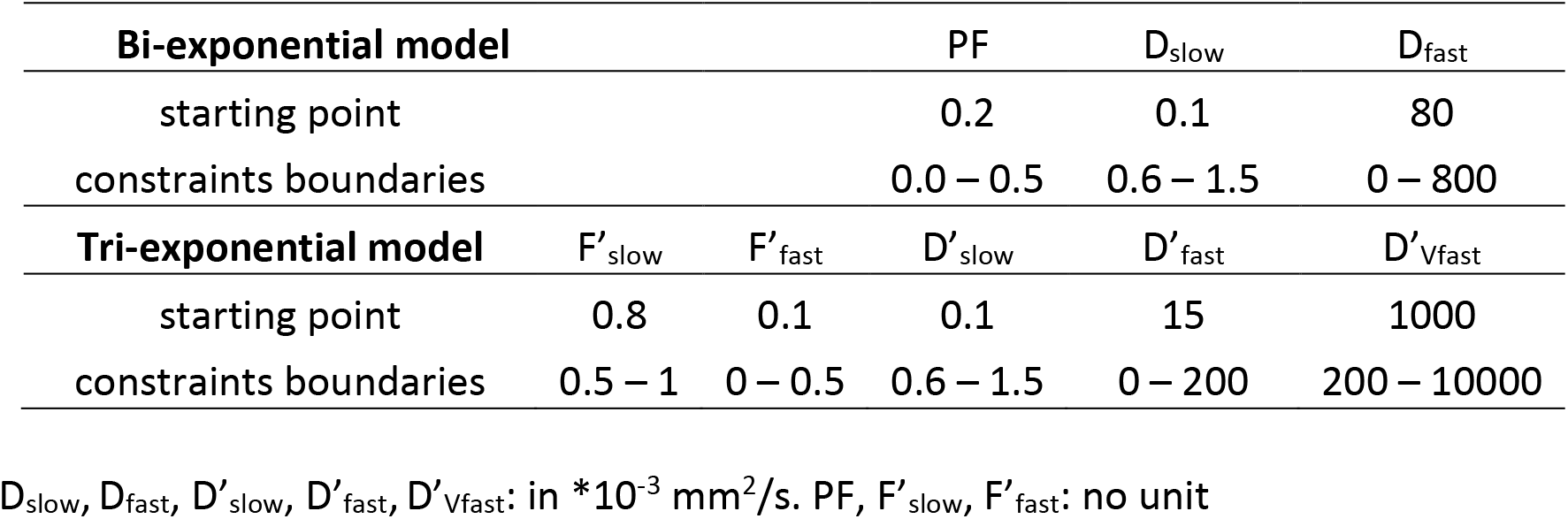
Starting points and constraints boundaries for fitting with Trust-Region Algorithm.

| Bi-exponential model |  |  | PF | D<sub>slow</sub> | D<sub>fast</sub> |
| --- | --- | --- | --- | --- | --- |
| starting point |  |  | 0.2 | 0.1 | 80 |
| constraints boundaries |  |  | 0.0 – 0.5 | 0.6 – 1.5 | 0 – 800 |
| Tri-exponential model | F' <sub>slow</sub> | F' <sub>fast</sub> | D' <sub>slow</sub> | D' <sub>fast</sub> | D' <sub>vfast</sub> |
| starting point | 0.8 | 0.1 | 0.1 | 15 | 1000 |
| constraints boundaries | 0.5 – 1 | 0 – 0.5 | 0.6 – 1.5 | 0 – 200 | 200 – 10000 |

$D_{\text{slow}}, D_{\text{fast}}, D'_{\text{slow}}, D'_{\text{fast}}, D'_{\text{vfast}}$ : in $\cdot 10^{-3} \text{ mm}^2/\text{s}$ . PF, $F'_{\text{slow}}, F'_{\text{fast}}$ : no unit

In addition to the IVIM parameter estimation based on individual scans, the measured signals at each *b*-value from the 50 scans were additionally averaged together, the averaged signals (total-averaged) were then fitted with the four approaches such as they were from a single scan (6, 14). The IVIM parameter values from pooled analysis are more likely to be close to physiological value (6). To assess which approach offers better fit, the residuals of the fit at each *b*-values, which was defined by the normalized measured signal intensity value minus the normalized predicted signal value with each fitting approach (SI_measured_ – SI_predicted_), were evaluated [Fig 2]. Data points relative to *b*=0 s/mm^2^ was considered to not be associated with any residual (equations [1], [2]). In addition, signal measurement is more reliable at *b*=0 s/mm^2^ as there is no diffusion gradient applied at this *b*-value.

**Figure 2.**
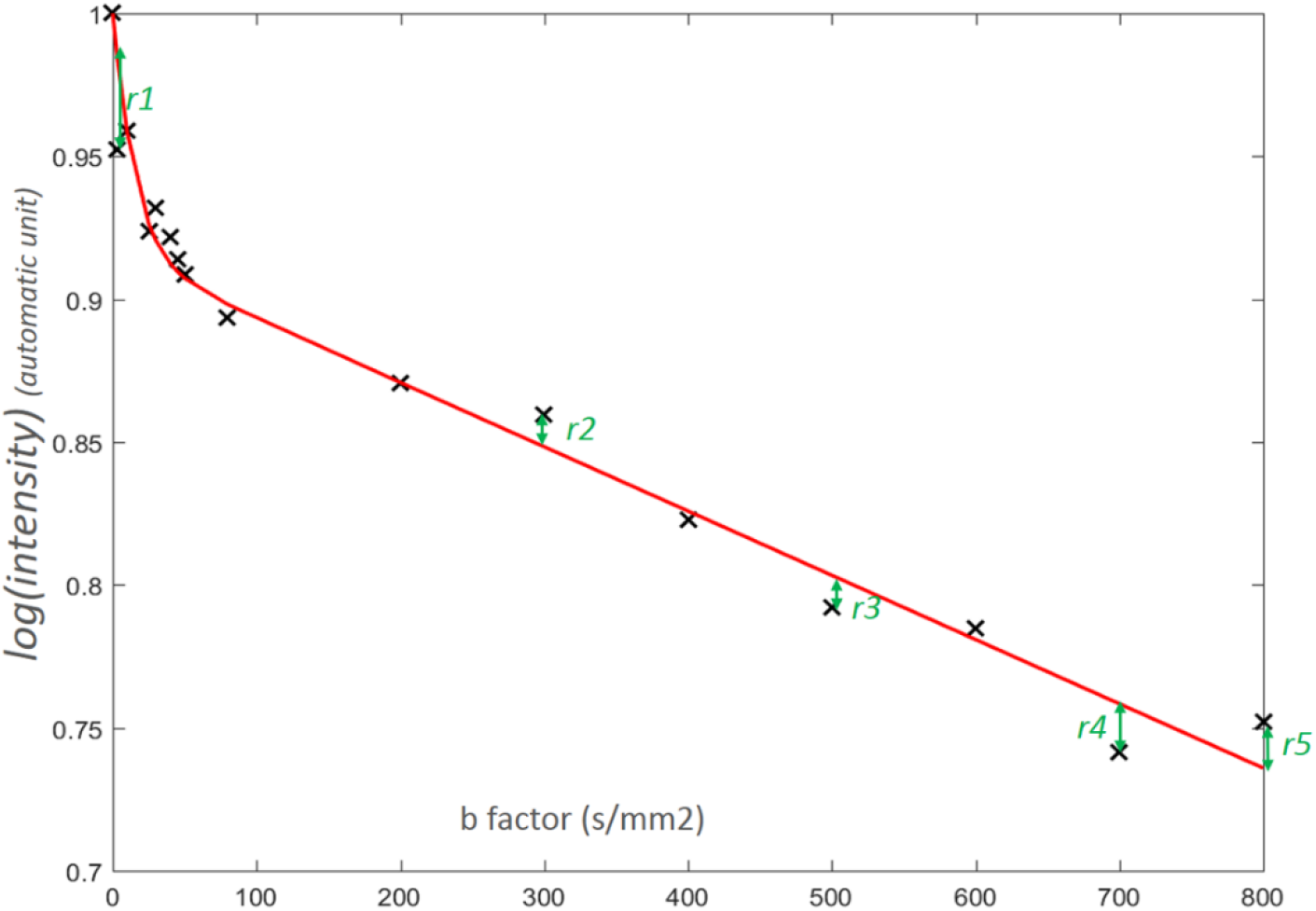
Graphic demonstration of definition of residual. Evidential residuals are shown at *b*=3 (*r1*), 300 (*r2*), 500 (*r3*), 700 (*r4*), and 800 (*r5*) s/mm^2^. *r1*=(signalmeasured) – (signalpredicted) at *b*=3 s/mm^2^, and thus gives a negative residual value, *r5*=(signalmeasured) – (signalpredicted) at *b*=800 s/mm^2^, and thus gives a positive residual value. Residuals at other *b*-values generally exist though with smaller values.

To assess which fitting method is favorable, in addition to visual graph assessment, extra sum-of-squares F-test (15, 16), Akaike’s information criterion (AIC), the second-order Akaike information criterion (AICc), delta AIC (ΔAIC), and evidence ratio were applied (16, 17) (supplement document 4). Intra-scan repeatability between scan 1.1 and scan 1.2, and inter-scan reproducibility between scan 1.1 and scan 2 of parameters were assessed by the average of coefficient of variation (CoV), and the within-subject coefficient of variation (_w_CoV), as previously described (4,5). Statistical analysis was performed using MedCalc Statistical Software (version 17.6, MedCalc Software bvba, Ostend, Belgium). SPSS software (IBM Corp. Released 2013. IBM SPSS Statistics for Windows, Version 22.0. Armonk, NY: IBM Corp) was used for the graphical analyzes.

## Results

Estimated IVIM parameters are shown in Table 2 (individual’s results in supplement tables 1 and 2). For bi-exponential model’s individual scans, D_slow_ values derived from full fitting were slightly higher than those derived from segmented fitting. The PF values derived from full fitting were lower than those derived from segmented fitting. D_fast_ values derived from full fitting were substantially higher than those derived from segmented fitting. For tri-exponential model’s individual scans, IVIM parameter were overall similar between full fitting and segmented fitting. The standard deviation of parameters related to perfusion (D_fast_, PF, D’_fast_, D’_Vfast_, F’_fast_, F’_Vfast_) were substantial, especially for D’_Vfast_ (Table 2). D’_Vfast_ value estimated with total-averaged data was lower than the mean of D’_Vfast_ value estimated by individual scan’s fits (table 2).

**Table 2.** IVIM parameters (mean± standard deviation) derived from bi-/tri-exponential models with full fitting or segmented fitting*.

| Bi-exponential | $D_{\text{slow}}$ | | $D_{\text{fast}}$ | PF | | | |
| --- | --- | --- | --- | --- | --- | --- | --- |
| Full fitting |  |  |  |  |  |  |  |
| Total-averaged (IC95%) | 1.14 (1.07 – 1.21) | - | 193.6 (99.7 – 287.5) | 16.9 (15.4 – 18.4) | - | - | - |
| All scans (n=50)** | 1.13 (±0.07) | - | 219.1 (±142.2) | 17.4 (±4.0) | - | - | - |
| Scans 1.1 (n=17) | 1.12 (±0.07) | - | 208.2 (±168.5) | 18.3 (±4.7) | - | - | - |
| Scans 1.2 (n=18) | 1.14 (±0.08) | - | 260.9 (±159.0) | 16.7 (±3.1) | - | - | - |
| Scans 2 (n=15) | 1.14 (±0.05) | - | 181.1 (±60.8) | 17.2 (±4.2) | - | - | - |
| Segmented fitting |  |  |  |  |  |  |  |
| Total-averaged (IC95%) | 1.03 (0.96 – 1.09) | - | 56.7 (38.7 – 74.8) | 21.3 (20.78 – 27.12) | - | - | - |
| All scans (n=50)** | 1.03 (±0.06) | - | 86.2 (±86.3) | 21.3 (±4.5) | - | - | - |
| Scans 1.1 (n=17) | 1.02 (±0.06) | - | 75.8 (±80.5) | 22.0 (±4.5) | - | - | - |
| Scans 1.2 (n=18) | 1.03 (±0.07) | - | 115.9 (±115.4) | 20.5 (±4.1) | - | - | - |
| Scans 2 (n=15) | 1.02 (±0.05) | - | 62.5 (±24.6) | 21.5 (±5.1) | - | - | - |
| Tri-exponential | $D'_{\text{slow}}^*$ | $D'_{\text{fast}}$ | $D'_{\text{vfast}}^*$ | PF' | $F'_{\text{slow}}$ | $F'_{\text{fast}}$ | $F'_{\text{vfast}}$ |
| Full fitting |  |  |  |  |  |  |  |
| Total-averaged (IC95%) | 0.98 (0.95 – 1.01) | 15.1 (11.9 – 18.3) | 445.0 (353.7 – 536.4) | 23.6 | 76.4 (75.2 – 77.6) | 11.8 (10.6 – 13.0) | 11.8 |
| All scans (n=50)** | 0.92 (±0.14) | 16.9 (±15.8) | 2132.8 (±3269.8) | 26.9 (±8.4) | 73.1 (±8.4) | 15.4 (±6.8) | 11.5 (±4.4) |
| Scans 1.1 (n=17) | 0.94 (±0.14) | 23.1 (±22.1) | 4173.8 (±4073.5) | 26.2 (±7.9) | 73.8 (±7.9) | 15.7 (±7.8) | 10.5 (±4.7) |
| Scans 1.2 (n=18) | 0.93 (±0.12) | 14.3 (±11.2) | 1579.0 (±2869.9) | 25.7 (±8.1) | 74.3 (±8.1) | 13.9 (±5.8) | 11.8 (±4.1) |
| Scans 2 (n=15) | 0.88 (±0.16) | 13.0 (±9.5) | 484.1 (±300.1) | 29.2 (±9.5) | 70.8 (±9.5) | 17.0 (±6.7) | 12.1 (±4.5) |
| Segmented fitting |  |  |  |  |  |  |  |
| Total-averaged (IC 95%) | 0.98 (0.95 – 1.02) | 15.4 (13.2 – 17.7) | 448.8 (361.7 – 535.8) | 23.4 | 76.6 (76.2 – 77.0) | 11.7 (10.9 – 12.5) | 11.7 |
| All scans (n=50)** | 0.98 (±0.07) | 16.1 (±7.8) | 1911.2 (±3190.5) | 23.6 (±4.9) | 76.4 (±4.9) | 11.9 (±4.0) | 11.7 (±4.1) |
| Scans 1.1 (n=17) | 0.98 (±0.06) | 18.3 (±9.1) | 3537.9 (±4173.4) | 24.0 (±4.7) | 76.0 (±4.7) | 12.7 (±4.6) | 11.4 (±5.0) |
| Scans 1.2 (n=18) | 0.99 (±0.07) | 13.9 (±4.4) | 1528.5 (±2889.5) | 22.9 (±4.5) | 77.1 (±4.5) | 11.0 (±3.7) | 11.9 (±3.3) |
| Scans 2 (n=15) | 0.98 (±0.07) | 16.0 (±9.2) | 527.0 (±358.6) | 23.8 (±5.7) | 76.2 (±5.7) | 12.1 (±3.7) | 11.7 (±4.2) |

\* Among the 50 scans, $D'_{\text{slow}}$ and $D'_{\text{vfast}}$ reached the lower fitting boundary of respectively 0.6 and 200 mm<sup>2</sup>/s for respectively four and one scans with full fitting. $D'_{\text{vfast}}$ reached the lower fitting bound of 200 mm<sup>2</sup>/s for three scans with segmented fitting (supplementary document 2). \*\* parameter mean ± SD of the 50 scans calculated one-scan by one-scan. $D_{\text{slow}}$ , $D_{\text{fast}}$ , $D'_{\text{slow}}$ , $D'_{\text{fast}}$ , and $D'_{\text{vfast}}$ are in 10<sup>-3</sup> mm<sup>2</sup>/s, PF, $F'_{\text{slow}}$ , $F'_{\text{fast}}$ , and $F'_{\text{vfast}}$ are in %. PF' calculated by (1 - $D'_{\text{slow}}$ ), $F'_{\text{vfast}}$ calculated by (1 - $F'_{\text{slow}}$ - $F'_{\text{fast}}$ ).

Graphical analysis of the fit residuals are shown in Fig. 3 and 4. Fig. 5 shows the curves of the total-averaged 50 scans signal using the four fitting approaches. Rather than randomness of the residuals, consistent trend of residuals with under- and over- estimation of the predicted signal value were observed with bi-exponential model. There was an inability of bi-exponential model to fit correctly very low (range: 3∼10 s/mm^2^) and low *b*-values (range: 25∼80 s/mm^2^) at the same time, with worse results using segmented fitting, which potentially led to inaccuracy in D_fast_ estimation. Segmented fitting especially overestimated the signal at *b*=3 and *b*=10 s/mm^2^(Fig. 3 (***a***) vs (***e***), Fig. 4 (***a***) vs (***e***)), however, it demonstrated lower error in signal prediction for higher *b*-values, leading to a better accuracy for D_slow_ estimation (Fig. 5, (***a***) vs (***b***)). Both tri-exponential model approaches showed relatively small errors in signal prediction which were more randomly and evenly distributed with slightly even smaller residuals for full fitting method (Fig. 3 (***i***) vs (***j***)). The R^2^ and adjusted-R^2^ were all higher than 0.95 for the four fitting approach, and adjusted-R^2^ favored tri-exponential model (supplement document 6). Extra sum-of-squares F-test and Akaike information criteria showed tri-exponential model was favored over bi-exponential model, and full fitting method was overall favored over segmented method (supplementary tables 3, 4). F-test using the fits of total-averaged signal also favored tri-exponential model, whatever the fitting methods (p<0.001 in all cases). For tri-exponential model, full fitting also offered even smaller residuals than segmented fitting, however, the difference was small (supplementary table 3, 4), which was consistent with results of Fig. 3, 4 and 5.

**Figure 3.**
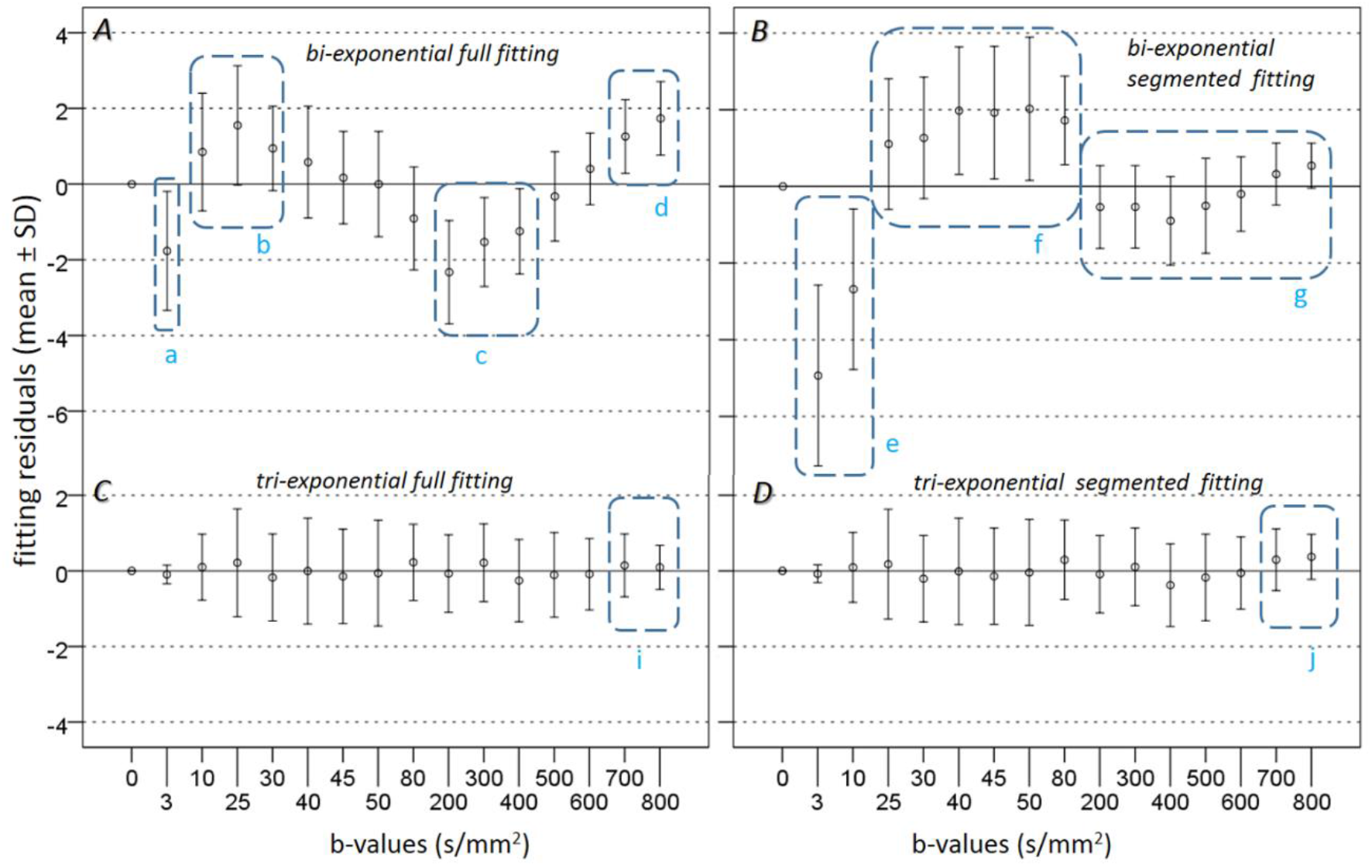
Tri-exponential fitting models show smaller and more randomly distributed residual compared with bi-exponential fitting models. Errors bars: mean ± SD (in %, n=50 scans). Bi-exponential full fitting (A) shows strong errors in predicted signal with negative residual for very low *b*-value (*b*=3 s/mm^2^) (***a***) and consecutive positive residuals for low *b*-values (***b***), leading potentially to an inaccuracy of D_fast_ estimation. For higher *b*-values, residuals are shown as negative (***c***) and then positive (***d***), leading potentially to inaccuracy of D_slow_ estimation. Bi-exponential segmented fitting (B) shows strong deviation with negative residuals for very low *b*-value (***e***), then consecutive positive residuals for low *b*-values (***f***), suggesting poor fitting for *b*-values lower than 200 s/mm^2^ and inaccuracy of Dfast estimation. For *b*-values ≥ 200 s/mm^2^, (B) shows the residuals are close to the reference line, consistent with the segmented method used (D_slow_ calculated first by linear regression with smaller residuals). Tri-exponential models (C, D) show smaller and more random distribution of the residuals, with segmented fitting showing slightly higher residuals at *b*=700 and 800 s/mm^2^(***i*** vs ***j***). Residuals significantly differ from 0 for *b*=3, 10, 25, 30, 40, 80, 200, 300, 400, 600, 700, 800 s/mm^2^ with bi-exponential full fitting (p<0.05, one sample t-test) and differ from 0 for all *b*-values (p<0.05) except *b*=600 s/mm^2^ with bi-exponential segmented fitting. With tri-exponential model, residuals differ significantly from 0 only for *b*=3 s/mm^2^ with full fitting method (p<0.05) and for *b*=3, 400, 700, 800 s/mm^2^ with segmented fitting method (p<0.05).

**Figure 4.**
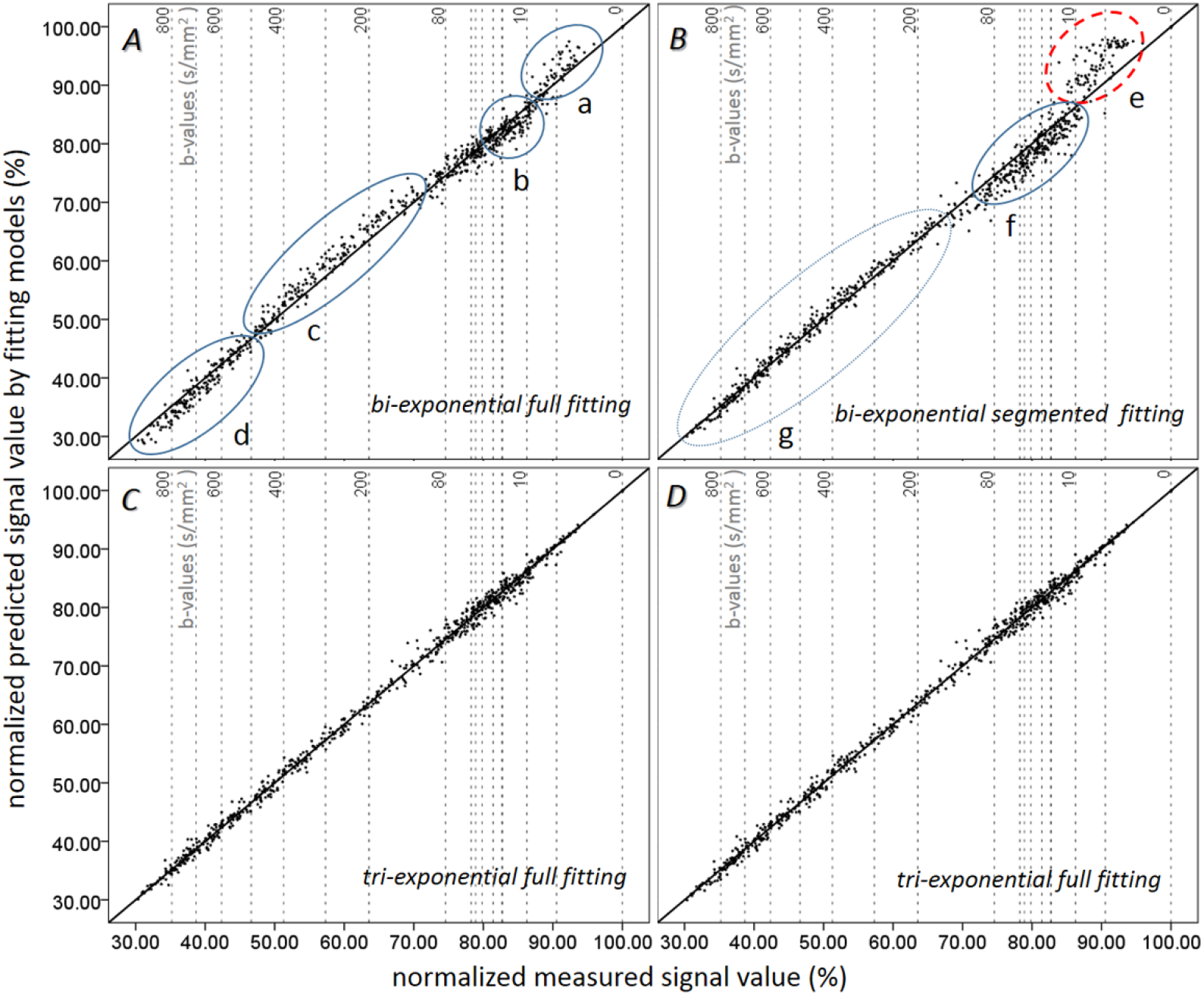
Tri-exponential fitting models result in a better prediction of the diffusion signal decay than bi-exponential fitting models. The graphs show the normalized predicted signal values vs. normalized measured signal values for all scans (n=50). The oblique lines show perfect predictions where the signal values derived from model fitting are equal to the measured signal values. For bi-exponential model, full fitting (A) shows overestimation of the predicted signal for very low *b*-values (***a***), and for *b*-values range 80 ∼400 s/mm^2^ (***c***); while at *b*-values range 10∼40 s/mm^2^ (***b***) and *b*-values range 600∼800 s/mm^2^ (***d***) the predicted signals are underestimated. The segmented fitting (B) shows a stronger overestimation of the predicted signal for very low *b*-values (***e***), and a trend of underestimation of the predicted signal for low *b*-values (***f***). On the other hand, for the high *b*-value part (≥ 200 s/mm^2^), the predicted signal values are more evenly distributed both below and above the reference line, i.e. the over- and under-estimations are more random and did not show an apparent trend (***g***). For tri-exponential model, both full fitting (C) and segmented fitting (D) show the predicted signals randomly distributed around the reference line with less deviation from measured signal values.

**Figure 5.**
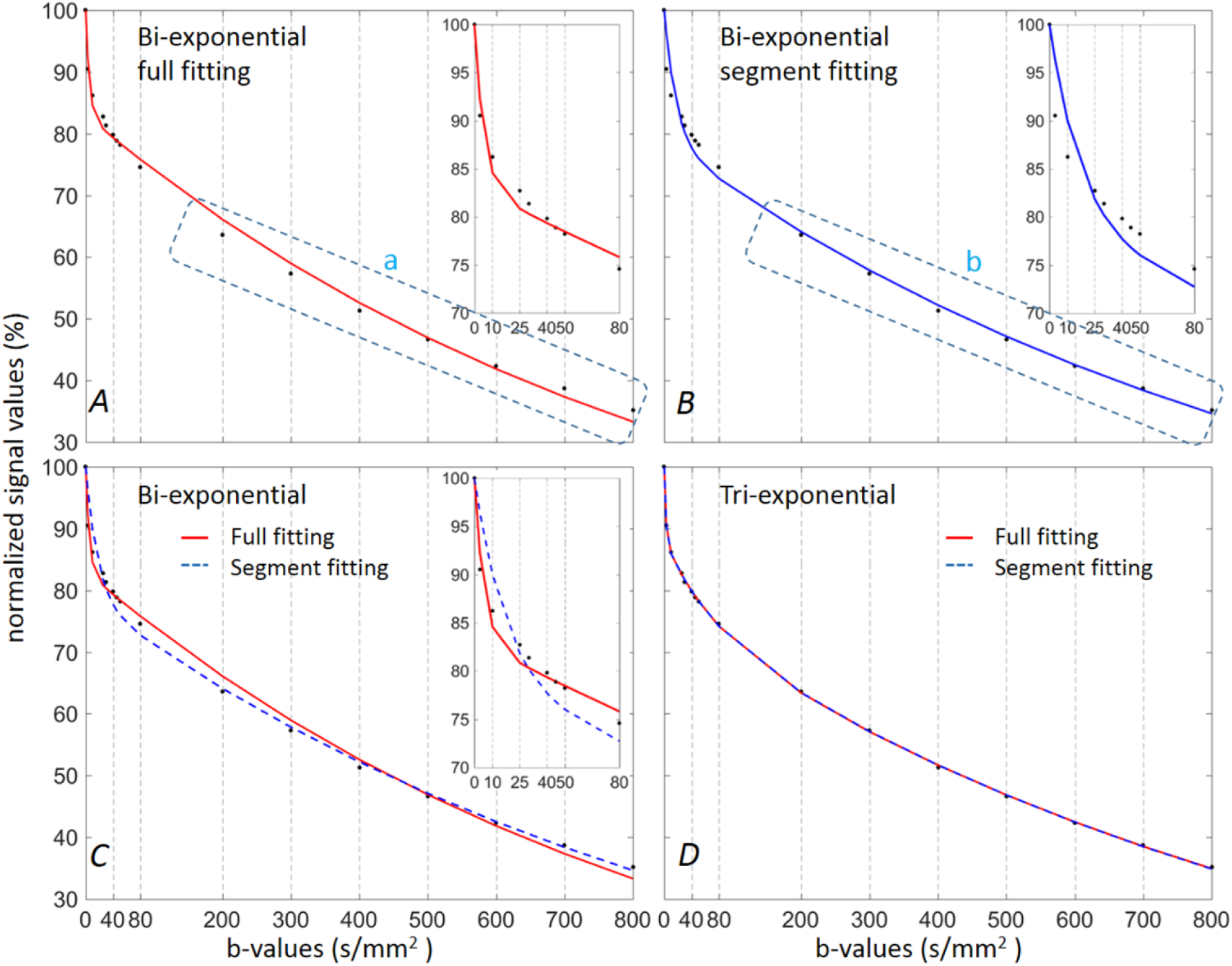
A comparison of total-averaged signal fitted curves using bi-/tri-exponential models with full or segmented fitting. For bi-exponential model, both fittings (A, B) do not fit well the initial part of the diffusion signal decay. Full fitting (A) tends to fit better at very low *b*-values (*b*=3, 10 s/mm^2^) and some low *b*-values (*b*=40, 45, 50 s/mm^2^) than segmented fitting model, and shows a steeper slope of the initial curve, leading to a higher D_fast_ value; while segmented fitting may underestimate Dfast (C). Segmented fitting model fits better at *b* ≥ 200 s/mm^2^, which leads to a better accuracy for D_slow_ estimation; while full fitting shows a slightly steeper slope leading to a higher D_slow_ value (***a*** vs ***b***). For the tri-exponential model, both fittings (D) show a good fit of diffusion signal decay, with the two fitted curves almost indistinguishable.

The best repeatability and reproducibility were found for D_slow_ with both fitting methods (Table 3). For bi-exponential model, D_fast_ was less repeatable/reproducible, being worse with full fitting method. For tri-exponential model, F’_slow_ and D’_slow_ showed the best repeatability/reproducibility results, with even better results with segmented fitting. For tri-exponential model, less repeatability and reproducibility were noted for perfusion related parameters (D’_fast_, D’_Vfast_, F’_fast_ and F’_Vfast_) with worse results for D’_Vfast_, with segmented fitting method achieving better repeatability/reproducibility than full fitting method (Table 3). Repeatability and reproducibility of PF’ calculated using (1- F’_slow_) of tri-exponential segmented fitting were comparable to PF estimated by bi-exponential model.

**Table 3.** Scan-rescan repeatability and reproducibility of IVIM parameters.

| Bi-exponential models |  | Full fitting |  | Segmented fitting |  |
| --- | --- | --- | --- | --- | --- |
| Parameters |  | Repeatability<br>(n=17) | Reproducibility<br>(n=14) | Repeatability<br>(n=17) | Reproducibility<br>(n=14) |
| D<sub>fast</sub> | Mean CoV (range) | 49.4 (4.3 – 110.0) | 43.9 (6.9 – 98.9) | 31.5 (0.2 – 111.4) | 22.6 (0.4 – 53.2) |
|  | Within-subject CoV | 57.3 | 57.3 | 67.3 | 28.3 |
| D<sub>slow</sub> | Mean CoV (range) | 2.2 (0.0 – 5.9) | 3.7 (0.1 – 10.0) | 3.5 (0.1 – 8.2) | 2.3 (0.7 – 5.3) |
|  | Within-subject CoV | 2.6 | 4.8 | 4.4 | 2.6 |
| PF | Mean CoV (range) | 13.5 (0.5 – 58.6) | 12.4 (3.4 – 32.7) | 14.1 (1.1 – 50.9) | 9.3 (0.1 – 28.1) |
|  | Within-subject CoV | 20.8 | 15.0 | 18.6 | 11.5 |
| Tri-exponential models |  | Full fitting |  | Segmented fitting |  |
| Parameters |  | Repeatability<br>(n=17) | Reproducibility<br>(n=14) | Repeatability<br>(n=17) | Reproducibility<br>(n=14) |
| D'<sub>slow</sub> | Mean CoV (range) | 10.4 (0.8 – 40.3) | 13.7 (1.4 – 38.6) | 4.3 (0.1 – 9.6) | 2.4 (0.2 – 6.2) |
|  | Within-subject CoV | 13.1 | 17.0 | 5.3 | 3.1 |
| D'<sub>fast</sub> | Mean CoV (range) | 48.7 (1.3 – 132.5) | 61.2 (2.6 – 131.6) | 28.5 (1.8 – 75.1) | 37.8 (2.6 – 97.4) |
|  | Within-subject CoV | 105.0 | 113.9 | 44.0 | 56.4 |
| D'<sub>vfast</sub> | Mean CoV (range) | 69.9 (0.1 – 134.7) | 86.6 (3.1 – 133.5) | 60.8 (1.3 – 130.6) | 75.8 (2.7 – 133.4) |
|  | Within-subject CoV | 132.0 | 165.4 | 139.6 | 179.1 |
| F'<sub>slow</sub> | Mean CoV (range) | 8.3 (1.0 – 27.0) | 9.3 (0.2 – 25.0) | 4.1 (0.1 – 11.7) | 3.0 (0.2 – 6.2) |
|  | Within-subject CoV | 10.1 | 11.5 | 5.2 | 3.7 |
| F'<sub>fast</sub> | Mean CoV (range) | 32.8 (1.1 – 57.9) | 29.8 (0.1 – 78.5) | 26.3 (0.3 – 59.2) | 21.6 (2.0 – 50.5) |
|  | Within-subject CoV | 39.7 | 41.6 | 32.5 | 28.2 |
| F'<sub>vfast</sub> | Mean CoV (range) | 35.8 (8.9 – 93.1) | 25.7 (0.1 – 89.0) | 26.2 (7.0 – 62.8) | 19.2 (2.7 – 33.5) |
|  | Within-subject CoV | 44.0 | 34.3 | 34.4 | 21.2 |
| PF' | Mean CoV (range) | 21.3 (3.4 – 49.7) | 21.4 (0.5 – 47.3) | 13.1 (0.3 – 43.6) | 9.7 (0.4 – 23.1) |
|  | Within-subject CoV | 28.4 | 29.9 | 16.9 | 11.4 |

CoV: coefficient of variation in %. Within-subject CoV: within-subject coefficient of variation in % (4, 5). PF' calculated by (1 - D'<sub>slow</sub>), F'<sub>vfast</sub> calculated by (1 - F'<sub>slow</sub> - F'<sub>fast</sub>).

For tri-exponential model, both full fitting and segmented fitting provided a good estimation of liver measured signal (Table 4). For the predicted signal, the very fast compartment (C’_Vfast_) contributed only for around 11.7-11.8% of the signal at *b*=0 s/mm^2^ and only around 3.4% of the signal at *b*=3 s/mm^2^, and its contribution was negligible at *b*-value ≥ 10 s/mm^2^. Contribution of the fast compartment (C’_fast_) was similar to C’_Vfast_ at *b*=0 s/mm^2^ and decreased slower, becoming negligible at *b*≥200 s/mm^2^. The slow compartment (C’_slow_) contributed to the majority of the total signal, starting at 76% at *b*=0 s/mm^2^, reaching almost 100% at b≥200 s/mm^2^. Considering all the *b*-values, C’_slow_, C’_fast_ and C’_Vfast_ contributions to the total DW signal measured were 92.1, 6.6, and 1.4% respectively. Bi-exponential model had a bigger difference between measured signal and predicted signal (Table 5). For the predicted signal, with bi-exponential full fitting the perfusion related diffusion compartment (C_fast_) contributed for around 16.9% of the total signal at *b*=0 s/mm^2^, then decreased to become negligible at b≥25 s/mm^2^, which is quite similar to the C’_Vfast_ contribution of the tri-exponential model. With bi-exponential segmented fitting, Cfast contributes around 21.4% at *b*=0 s/mm^2^, which was closer to the sum of the contributions of two compartments related to perfusion of the tri-exponential model (i.e. C’_fast_ and C’_Vfast_). The contribution became negligible at *b*≥80 s/mm^2^ which was consistent with the segmented fitting method and the chosen threshold for D_slow_ calculation. This agreed with the observation in Table 2 that D_fast_ in bi-exponential full fitting had much higher value (close to C’_Vfast_ contribution) and was less stable in reproducibility (Table 3).

**Table 4.** Signal contribution of the slow, fast, and very fast compartments using tri-exponential model and total-averaged 50 scans at each *b*-value.

| | Average measured SI (±SD) | Predicted signal by full fitting | | | | Predicted signal by segmented fitting | | | |
| --- | --- | --- | --- | --- | --- | --- | --- | --- | --- |
| $b$-value (s/mm<sup>2</sup>) | | TSI | $C'_{\text{slow}}/\text{TSI}$ | $C'_{\text{fast}}/\text{TSI}$ | $C'_{\text{vfast}}/\text{TSI}$ | TSI | $C'_{\text{slow}}/\text{TSI}$ | $C'_{\text{fast}}/\text{TSI}$ | $C'_{\text{vfast}}/\text{TSI}$ |
| 0 | 100.0 ± 0.0 | 100.0 | 76.4 | 11.82 | 11.78 | 100.0 | 76.6 | 11.66 | 11.72 |
| 3 | 90.5 ± 2.7 | 90.6 | 84.1 | 12.47 | 3.42 | 90.6 | 84.3 | 12.30 | 3.37 |
| 10 | 86.3 ± 3.3 | 86.0 | 88.0 | 11.83 | 0.16 | 86.0 | 88.2 | 11.62 | 0.15 |
| 25 | 82.7 ± 4.2 | 82.7 | 90.2 | 9.81 | 0.00 | 82.7 | 90.4 | 9.59 | 0.00 |
| 30 | 81.4 ± 4.0 | 81.7 | 90.8 | 9.21 | 0.00 | 81.7 | 91.0 | 8.98 | 0.00 |
| 40 | 79.8 ± 3.5 | 79.9 | 91.9 | 8.09 | 0.00 | 79.9 | 92.1 | 7.86 | 0.00 |
| 45 | 78.9 ± 3.9 | 79.1 | 92.4 | 7.59 | 0.00 | 79.1 | 92.6 | 7.36 | 0.00 |
| 50 | 78.2 ± 4.0 | 78.3 | 92.9 | 7.11 | 0.00 | 78.3 | 93.1 | 6.88 | 0.00 |
| 80 | 74.6 ± 4.1 | 74.2 | 95.2 | 4.77 | 0.00 | 74.2 | 95.4 | 4.57 | 0.00 |
| 200 | 63.6 ± 3.9 | 63.4 | 99.1 | 0.92 | 0.00 | 63.5 | 99.2 | 0.84 | 0.00 |
| 300 | 57.3 ± 3.6 | 57.1 | 99.8 | 0.23 | 0.00 | 57.2 | 99.8 | 0.20 | 0.00 |
| 400 | 51.3 ± 3.0 | 51.7 | 99.9 | 0.06 | 0.00 | 51.7 | 100.0 | 0.05 | 0.00 |
| 500 | 46.6 ± 3.4 | 46.9 | 100.0 | 0.01 | 0.00 | 46.9 | 100.0 | 0.01 | 0.00 |
| 600 | 42.3 ± 2.9 | 42.5 | 100.0 | 0.00 | 0.00 | 42.5 | 100.0 | 0.00 | 0.00 |
| 700 | 38.7 ± 2.7 | 38.5 | 100.0 | 0.00 | 0.00 | 38.5 | 100.0 | 0.00 | 0.00 |
| 800 | 35.2 ± 2.4 | 34.9 | 100.0 | 0.00 | 0.00 | 34.9 | 100.0 | 0.00 | 0.00 |

Average measured signal intensity (SI): average of the normalized measured liver signal of the 50 selected scans. $C'_{\text{slow}}$ , $C'_{\text{fast}}$ , $C'_{\text{vfast}}$ , are related to the first (slow) compartment signal [ $F'_{\text{slow}} \times \exp(-b \times D'_{\text{slow}})$ ], the second (fast) compartment signal [ $F'_{\text{fast}} \times \exp(-b \times D'_{\text{fast}})$ ], and the third (very fast) compartment signal [ $F'_{\text{vfast}} \times \exp(-b \times D'_{\text{vfast}})$ ] respectively. TSI: Total diffusion weighted imaging signal predicted by the model (sum of the tri-compartment signals). $C_i/\text{TSI}$ : signal contribution of the compartment $i$ in the total predicted signal, in %.

**Table 5.** Signal contribution of the slow and fast compartments using bi-exponential models and total-averaged 50 scans at each *b*-values.

| | Average measured SI (±SD) | Predicted signal by full Fitting | | | Predicted signal by Segmented Fitting | | |
| --- | --- | --- | --- | --- | --- | --- | --- |
| $b$-value (s/mm<sup>2</sup>) | | TSI | C<sub>slow</sub>/TSI | C<sub>fast</sub>/TSI | TSI | C<sub>slow</sub>/TSI | C<sub>fast</sub>/TSI |
| 0 | 100.0 ± 0.0 | 100.0 | 83.1 | 16.92 | 100.0 | 78.6 | 21.42 |
| 3 | 90.5 ± 2.7 | 92.3 | 89.7 | 10.26 | 96.4 | 81.3 | 18.74 |
| 10 | 86.3 ± 3.3 | 84.6 | 97.1 | 2.89 | 89.9 | 86.5 | 13.51 |
| 25 | 82.7 ± 4.2 | 80.9 | 99.8 | 0.17 | 81.8 | 93.7 | 6.34 |
| 30 | 81.4 ± 4.0 | 80.3 | 99.9 | 0.06 | 80.1 | 95.1 | 4.88 |
| 40 | 79.8 ± 3.5 | 79.4 | 100.0 | 0.01 | 77.6 | 97.1 | 2.85 |
| 45 | 78.9 ± 3.9 | 78.9 | 100.0 | 0.00 | 76.7 | 97.8 | 2.17 |
| 50 | 78.2 ± 4.0 | 78.5 | 100.0 | 0.00 | 75.9 | 98.3 | 1.65 |
| 80 | 74.6 ± 4.1 | 75.8 | 100.0 | 0.00 | 72.6 | 99.7 | 0.32 |
| 200 | 63.6 ± 3.9 | 66.1 | 100.0 | 0.00 | 64.0 | 100.0 | 0.00 |
| 300 | 57.3 ± 3.6 | 59.0 | 100.0 | 0.00 | 57.8 | 100.0 | 0.00 |
| 400 | 51.3 ± 3.0 | 52.6 | 100.0 | 0.00 | 52.1 | 100.0 | 0.00 |
| 500 | 46.6 ± 3.4 | 46.9 | 100.0 | 0.00 | 47.1 | 100.0 | 0.00 |
| 600 | 42.3 ± 2.9 | 41.9 | 100.0 | 0.00 | 42.5 | 100.0 | 0.00 |
| 700 | 38.7 ± 2.7 | 37.3 | 100.0 | 0.00 | 38.3 | 100.0 | 0.00 |
| 800 | 35.2 ± 2.4 | 33.3 | 100.0 | 0.00 | 34.6 | 100.0 | 0.00 |

Average measured signal intensity (SI): average of the normalized measured liver signal of the 50 selected scans. TSI: Total diffusion weighted imaging signal predicted by the model. C<sub>slow</sub> and C<sub>fast</sub>, are related to the true diffusion compartment signal $[(1 - PF) \times \exp(-b \times D_{\text{slow}})]$ and the pseudo-diffusion compartment signal $[PF \times \exp(-b \times D_{\text{fast}})]$ respectively. Si/TSI: Signal contribution of the compartment $i$ in the total signal predicted signal, in %.

## Discussion

This study demonstrates that, for the fitting of diffusion MR signal decay in healthy livers, tri-exponential model provides overall better fit compared with bi-exponential model. With tri-exponential model, the actually measured signal values and predicted signal values show smaller and more randomly distributed residual errors than with bi-exponential model. Analysis of F-test and Akaike information criteria also favored the tri-exponential model, meaning the addition of new parameters was justified. The better fitting results of the tri-exponential model can be due to the presence of an additional very fast perfusion-related compartment, consistent with the observed very fast decay of the DW imaging signal for very low *b*-value not described in the original IVIM theory (1, 6, 7). According to Cercueil *et al*. (6) and Gurney-Champion *et al*. (18), this very fast signal decay (D’_Vfast_ and F’_Vfast_) could be explained by vessel components included in the ROIs (6, 18). Our study showed a slow (D’_slow_=0.98×10^-3^ mm^2^/s; F’_slow_=76.4 (with full fitting) or 76.6% (with segmented fitting)), a fast (D’_fast_=15.1 or 15.4×10^-3^ mm^2^/s; F’_fast_=11.8 or 11.7%) and a very fast (D’_Vfast_=445.0 or 448.8×10^-3^ mm^2^/s; F’_Vfast_=11.8 or 11.7 %) diffusion compartments. Cercueil *et al*. reported tri-exponential IVIM model determined three diffusion compartments with full fitting: a slow (D’_slow_=1.35×10^-3^ mm^2^/s; F’_slow_=72.7%), a fast (D’_fast_=26.50×10^-3^ mm^2^/s; F’_fast_=13.7%) and a very fast (D’_Vfast_=404.00×10^-3^ mm^2^/s; F’_Vfast_=13.5%) diffusion compartment (6). The F’_slow_, F’_fast_, and F’_Vfast_ values are similar between our results and Cercueil *et al*.’s results, while D’_slow_ and D’_fast_ values are slightly different. These differences may be primarily due to the fact that different *b*-value distributions were used in these two studies. Note that Cercueil *et al*.’s results were also different between pilot and validation studies despite the same hardware, suggesting *b*-value distribution influenced the parameters values.

Our study showed bi-exponential model did not perform well in fitting very low and low *b*-values at the same time, which led to inaccuracy in D_fast_ estimation; and errors in fitting were even more substantial with segmented fitting (Fig 3, 4). For bi-exponential model, the segmented fitting promoted the accuracy of Dslow estimation at the expense of accuracy in D_fast_ estimation. D_slow_ repeatability/reproducibility were similar between full fitting and segmented fitting, but the fit for high *b*-values was better with segmented fitting. In addition, the repeatability/reproducibility of PF and D_fast_ were better with segmented fitting since less parameters are estimated by non-linear regression (i.e. only D_fast_ was estimated instead of the three parameters). Bi-exponential full fitting fitted slightly worse at high *b*-values, with a steeper slope, meaning Dslow could be over-estimated (Fig. 3, 4 and 5, Table 2). Therefore, segmented fitting is probably more applicable for studying disease processes which primarily change the slow diffusion compartment. On the other hand, for assessing changes in microcirculation (pseudo-diffusion compartment), full fitting method is more appropriate (Fig. 3 (***a***) vs (***e***), and Fig. 4 (***a***) vs (***e***)). However, at very low and low *b*-values, for both full fitting and segmented fitting, bi-exponential model did not fit as well as tri-exponential model, while IVIM parameter associated with these *b*-values are essential to assess the disease changes associated with fast diffusion (perfusion related diffusion). In fact, many liver pathologies may initially affect fast diffusion (19, 20). Recent literature review also showed that among nine patient studies, only D_fast_, despite being the least stable, consistently demonstrated liver fibrosis is progressively associated with a reduced measurement (3). D_fast_ estimation depends on the number and distribution of low *b*-values. D_fast_ tends to be under-estimated when few low *b*-values are included (3, 21), while it tends to increase when more very low *b*-values are included, as very low *b*-value are essential to catch the initial fast decay of DW imaging (3, 6) (supplementary Fig. 1, 2).

The reproducibility of bi-exponential models in this study is consistent with recent reports (4, 5, 18, 20), while this study is the first to explore tri-exponential model IVIM parameter reproducibility with two perfusion related components. For bi-exponential model, D_slow_ was most stable, followed by PF, and D_fast_ was less reproducible. A recent study suggested that reproducibility of IVIM parameters could be improved by a ‘data cleaning process’ which was partially used in this study (4). This current study showed bi-exponential segmented fitting tended to have these parameters more stabilized, but at the cost of stronger error in predicted signal for low *b*-values. In the same way as bi-exponential model, tri-exponential model derived good reproducibility for parameters related to the true diffusion, such as D’_slow_ and F’_slow_, while less stable for parameters related to perfusion, such as D’_fast_, D’_Vfast_, F’_fast_, and F’_Vfast_. Theoretically, the more coefficients are added to a model, the more likely estimated parameters become less stable. That D’_Vfast_ showed apparent instability could be also partially explained by the *b*-values distribution in this study. *b*-values distribution in this study was still not optimized for a tri-exponential model, with under-sampling of diffusion signal at very low *b*-values. A better stability for D’_Vfast_ can be achieved by adding more very low *b*-values (authors’ unpublished results). However, parameters related to extra-cellular water molecules motion, D’_slow_ and F’_slow_, showed a better reproducibility, globally similar to D_slow_ and PF reproducibility of bi-exponential segmented method model. These observations support the signal decay analysis where for high *b*-values the signal decay depends only on the slow decay compartment (table 4, 5).

For very low *b*-values (*b*-values ≤ 10 s/mm^2^), all compartments contributes to the DW imaging signal. However, the contribution of the third compartment was low at *b*=3 s/mm^2^(≈3.4%) and negligible at *b*=10 s/mm^2^ (≈0.15%) (Table 4). D’_Vfast_ estimated with total-averaged data was substantially lower than D’_Vfast_ estimated with the average of D’_Vfast_ values estimated by individual scan fits. However, the D’_Vfast_ value estimated in this study (445.0 ×10^-3^ mm^2^/s with full fitting and 448.8×10^-3^ mm^2^/s with segmented fitting) is quite similar to Cercueil *et al*.’s report (404.00×10^-3^ mm^2^/s). It can be hypothesized that averaging the data of multiple scans leads to a better estimation of the “true” D’_Vfast_ value. With total-averaged method, estimated C’_Vfast_ contribution indeed represented only 1.4% of the total DW measured signal considering all *b*-values (vs. 92% for C’_slow_). It highlights the limitation of accurate D’_Vfast_ value estimation in this context of low SNR imaging, especially using individual subject data. As previously noticed, this compartment was insufficiently sampled in our study. Interestingly, in a previous study where the lowest *b*-value was *b*=10 s/mm^2^ therefore reducing the tri-exponential to a bi-exponential model (13), D_fast_ value was 12.34± 3.06 ×10^−3^ mm^2^/s, thus close to our D’_fast_ value (≈15 ×10^−3^ mm^2^/s). In addition, that study showed a combined analysis of D_slow_, PF, and D_fast_ estimated from bi-exponential model allows separation of early stage of liver fibrosis from healthy livers (13). This may suggest that a precise estimation of the very fast compartment (D’_Vfast_ and F’_Vfast_) may not be essential in some clinical applications.

One limitation of this study is that it was performed on healthy livers. The adequacy of the tri-exponential model in diseased settings needs to be explored in further studies. In addition, we did not use *b*-values higher than 800 s/mm^2^. A faster component and a slower component of the true diffusion have been evaluated at high *b*-value (22, 23). However, in liver the signal intensities with *b*-values of more than 800 s/mm^2^ may be close to the noise level under many clinically practicable scan settings. As noted previously, we may have not sampled sufficient very low *b*-value for optimal C’_Vfast_ compartment fitting. Finally, we did not control the volunteers’ fasting state, while the hepatic flow may vary depending on the fasted/prandial status, therefor control the volunteers’ fasting state may further improve scan-rescan repeatability/reproducibility (24).

In conclusion, tri-exponential model provides a better fit for IVIM signal decay in the healthy liver than the classical bi-exponential model across the 0–800 s/mm^2^ range. For bi-exponential model, full fitting may be preferred for a better estimation for D_fast_ than segmented fitting, while segmented fitting offers better estimation of D_slow_. For tri-exponential model, the difference between full fitting and segmented fitting tends to be small concerning the fitting quality, but segmented fitting is preferred due to its better scan-scan reproducibility. The choice of signal decay model and fitting model may depend on the pathologies to be studied, and multiple model analysis can be applied for the same pathology. Our study represents the preliminary promising results of IVIM multiple compartments analysis, further development with better sampling with more very low and low *b*-values, motion correction, and better gradient coil can offer more precise analysis of IVIM parameters.

## Acknowledgement

Dr Olivier Chevallier was supported by a grant provided by the Société Française de Radiologie (SFR) together with the Collège des Enseignants de Radiologie de France (CERF). We thank Dr Weibo Chen, Philips Healthcare Shanghai, China, for setting-up the liver IVIM diffusion MRI acquisition protocol in Nanjing; and Mr Richard Yan Chak Li, research student at the Hong Kong University of Science and Technology, for discussions on Matlab programming.

## Supplementary Documents

### 1. Suppl Doc 1: The choice of threshold b-values for segmented fitting

The threshold *b*-value was chosen to be 80 s/mm^2^ for bi-exponential model in this study. The optimal threshold b-value remains controversial in literature. The most commonly thresholds used have been ≥ 100 s/mm^2^(1, 2) or ≥ 200 s/mm^2^(3–12). Since D_fast_ was significantly greater than D_slow_, its influence on signal decay was considered negligible for b-value higher than 100 – 200 s/mm^2^. The use of a lower threshold *b*-value (20 – 40 s/mm^2^) has also been suggested, as lower threshold value can lead to a minimization of residual sum of squares of the fit, thus smaller fitting error in signal prediction (13). We used *b*=80 s/mm^2^ as threshold value because this threshold b-value has been shown to offer better detection for early stage liver fibrosis (14). Recently it has also been shown that this threshold improves scan-rescan reproducibly (15). According to Cercueil *et al*. (16), the contribution of the pseudo-diffusion compartment at *b*=80 s/mm^2^ is low, between 5.08% for *b*=50 s/mm^2^ and 1.49% for *b*=100 s/mm^2^ of the total MR diffusion signal intensity, and therefore the contamination by perfusion element for D_slow_ estimation can be considered to be low for *b* ≥80 s/mm^2^. We consider this threshold would represent a good compromise between compartments separation, fit performance, reproducibility, and liver fibrosis detection performance.

For tri-exponential model, threshold *b*-value of *b*=200 s/mm^2^ was chosen in order to include more b-values for fast and very fast diffusion decays assessment, since we wanted to study and separate the two components of perfusion related diffusion. In addition, the contribution of the sum of both perfusion related compartments at *b*=200 s/mm^2^ was estimated at 0.12% of the total MR diffusion signal intensity, leading thus to a negligible contamination by the perfusion components for true diffusion compartment assessment and a lower loss of information for the fast components analysis (16).

Since in this study we are primarily concerned with the comparisons of bi-exponential model vs. tri-exponential model and full fitting vs. segmented fitting, different threshold *b*-values for bi-exponential and tri-exponential models can be chosen.

### 2. Suppl Doc 2:. Starting points and constraints boundaries for fitting with Trust-Region Algorithm (table 1 in the method section)

For bi-exponential model, starting points and constraints boundaries were chosen according to a similar study and studies results reported in a recent literature review (15, 17). However, for D_fast_ we chose a higher boundary value, since D_fast_ value tended to increase with the increase of the very low b-values number (5,16,17). For tri-exponential model, since D’_slow_ and F’_slow_ are theoretically similar to respectively D_slow_ and (1-PF) of bi-exponential model, for these two parameters we used the same starting points and boundaries as those in bi-exponential models. The starting points of F’_fast_, D’_fast_ and D’_Vfast_ were initially chosen according to Cercueil et al’s (16). However, after several trials using our data series, we found a trend to obtain lower value for F’_fast_, and D’_fast_, and higher value for D’_Vfast_. We thus changed the starting points according to our preliminary data results. Because the true values of these parameters are known to be unstable and have not yet been clearly defined in the literature (16), we decided to use wide boundaries, especially for D’_Vfast_.

All data were then processed using these fixed starting points and boundaries. Among the 50 scans, D’_slow_ and D’_Vfast_ reached the lower fitting boundary of respectively 0.6 and 200 mm^2^/s for respectively four and one scans with full fitting. D’_Vfast_ reached the lower fitting boundary of 200 mm^2^/s for three scans with segmented fitting. After reprocessing these data with lower boundary=0, the lowest D’_slow_ value was 0.49*10^-3^ mm^2^/s and the lowest D’_Vfast_ value was 88.2*10^-3^ mm^2^/s (thus not widely different from the lower boundaries).

### 3. Suppl Doc 3: Adjusted-R^2^ principle. (18)

The well-known coefficient of determination R^2^ allows a quantification of the goodness of the fit, evaluating the dispersion of the data points around the fitted curve, with a value between 0 and 1. The closer to 1 is the value, the smaller is the difference between the fitted values and the measured values. However, the addition of new parameters always increases the R^2^ that can be misleading. The adjusted-R^2^ allows a correction of this problem. Adjusted-R^2^ adjusts for the number of terms (parameters) in a model and is therefore a better indicator when comparing nested models (explained in Suppl Doc 4). If useful coefficient are added to the equation, the adjusted-R^2^ will increase. On the contrary if useless variables are added to a model, adjusted-R^2^ will decrease. Adjusted-R^2^ is always lower than or equals to R^2^. Different formulas exist for calculation of adjusted-R^2^ (19). In our article, we chose to calculate adjusted-R^2^ using the most commonly used formula (The Wherry formula-1): 

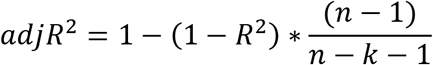

where n is the number of data point in the data sample (i.e. number of *b*-values in this study), k is the number of parameters.

However, a high R^2^ or adjusted-R^2^ does not always indicate a good model. A biased model with systematic over or under-estimation of the predicted value can present a higher R^2^ than an unbiased model with residuals more randomly distributed around 0. That is the reason why we also examined the residual plots of each fitting model.

### 4. Suppl Doc 4: Extra sum of squares F-test, Akaike’s Information Criterion (AIC), and evidence ratio

*Extra sum-of-squares F-test* is a statistical test used to compare nested models, which means that one model is a simpler case of the other. It tests the null hypothesis that the simpler model is preferable (20). Two models can be considered nested when the more complex model results from the addition of new parameter(s) to the simpler model. In our study, for the tri-exponential model we add two parameters to the bi-exponential model IVIM equation: D’_Vfast_ and F’_Vfast_. The models with more parameters in its equation tend to fit the data better (with smaller residual errors) than the model with fewer parameters, because the addition of parameters gives more flexibility to the fitting, resulting in smaller residual sum of square.

The *extra sum–of-squares F-test* is based on the difference in residual sum of squares of the models. However, the number of parameters of each models as well as the number of data points (number of *b*-value) are also taken into account. An F-ratio (F) is computed using the following equation:

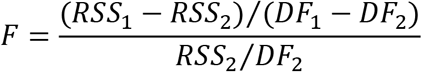

Where numbers 1 and 2 refer to the simpler (bi-exponential) and more complex (tri-exponential) models, respectively; RSS denotes the residual sum of squares; DF corresponds to the degree of freedom, which is equal to the number of data points (number of *b*-value) minus the number of parameters fitted by a model. Based on an F distribution, a p-value can be calculated from the F-ratio and the two DF values. In our study, if the F-ratio is associated with a p-value lower than 0.05, the more complex model (tri-exponential) is considered preferable than the simpler model (bi-exponential).

Akaike’s Information Criterion (AIC) permits to determine which model is more likely to be correct and quantify how much more likely (20, 21). Contrary to F-test, both nested and non-nested models can be compared by using AIC. AIC is calculated by the equation:

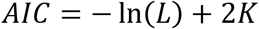

Where L is the maximum likelihood function, K is the number of parameters plus one. In our article, we use this last fundamental equation to compute AIC. For least square with normal distribution of residuals, AIC can be expressed as:

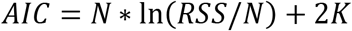

Where N is the number of data points (number of *b*-value). AIC value can be positive or negative, with no meaning of the sign itself.

The second order (or corrected) AICc is more accurate when N is small relative to K (ratio n/K < 40) and can be calculated by the equation:

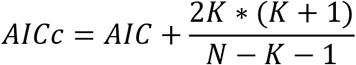

AIC and AICc values have no particular meaning and are not interpretable alone. Only the difference in AIC (or AICc) values is informative when comparing models. When comparing two models, the model presenting the lower AIC is more likely to be correct than the one with the higher AIC. The difference in AIC can also be calculated:

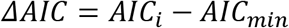

Where AIC_min_ corresponds to AIC value of the best model in the setting with the lowest AIC and AIC_i_ to the AIC of the model i. ΔAIC allows a quick comparison by quantifying the information loss when a model is used rather than the best approximating model. The larger is the ΔAIC, the less plausible is the model i as being the best approximating model. Classically, in comparison with a reference model (with lower AIC), models presenting ΔAIC ≤ 2, 4 ≤ ΔAIC ≤ 7 and ΔAIC ≥ 10, have respectively substantial support (evidence), considerable less support, and essentially no support.

The evidence ratio (or relative likelihood) permits to know how many times the model with the lowest AICc (model A) is more likely to be correct than the model with the highest AICc score (model B), and can be calculated by:

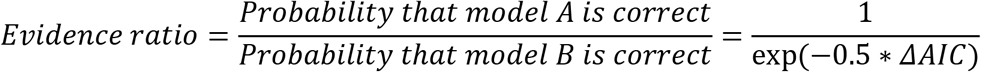

### 5. Suppl Doc 5: Comparison of R^2^ and adjusted-R^2^, Residual Sum of Squares, of tri-/bi-exponential models and full and segmented fitting methods with the individual scans’ data (n=50)

| Models and method | Mean $R^2$ | Min $R^2$ | Mean Adjusted-$R^2$ | Min Adjusted-$R^2$ | RSS (range) |
| --- | --- | --- | --- | --- | --- |
| Tri full | 99.72 | 99.02 | 99.59 | 98.54 | 17.1 (2.4 – 55.8) |
| Tri Seg | 99.71 | 99.02 | 99.57 | 98.54 | 17.8 (2.4 – 55.9) |
| Bi Full | 99.26 | 97.92 | 99.07 | 97.40 | 46.8 (11.7 – 119.1) |
| Bi Seg | 98.66 | 96.19 | 98.32 | 95.24 | 83.5 (16.9 – 218.0) |

Tri full: Tri-exponential full fitting, Tri seg: Tri-exponential segment fitting, Bi full: Bi-exponential full fitting, Bi full: Bi-exponential segment fitting. R^2^ and Adjusted-R^2^ in 10^-2^, no unit. RSS: residual sum of squares. Mean R^2^ / adjusted-R^2^ correspond to the mean R^2^ / adjusted-R^2^ value of the individual scans’ data fits. Min R^2^ / adjusted-R^2^ correspond to the minimum R^2^ / adjusted-R^2^ value of the individual scans’ data fits. R^2^ of tri-exponential model is higher than R^2^ of bi-exponential model, whatever the fitting method, except in one case where bi-exponential full fitting achieves better than tri-exponential segmented fitting. Tri full achieves higher adjusted-R^2^ than Tri seg, Bi full, Bi seg, in respectively 48 scans, 46 scans and 49 scans. Tri full achieves higher adjusted-R^2^ than Bi full and Bi seg in respectively 47 and 49 scans. Bi full achieves higher adjusted-R^2^ than Bi seg in all 50 scans.

---

**Supplementary Table 1.** IVIM parameters estimated with bi-exponential model (Vol.: Volunteers; D_slow_, Dfast in 10^-3^ mm^2^/s; PF: %).

| Vol. | Scan session | Full fitting |  |  | Segment fitting |  |  |
| --- | --- | --- | --- | --- | --- | --- | --- |
| | | $D_{\text{slow}}$ | $D_{\text{fast}}$ | PF | $D_{\text{slow}}$ | $D_{\text{fast}}$ | PF |
| 1 | 1.1 | 1.06 | 91.9 | 18.4 | 1.00 | 58.53 | 20.6 |
|  | 1.2 | 1.08 | 378.7 | 18.8 | 0.95 | 75.96 | 23.5 |
|  | 2 | 1.22 | 281.8 | 16.6 | 1.04 | 44.06 | 22.9 |
| 2 | 1.1 | 1.25 | 157.7 | 13.6 | 1.18 | 65.67 | 16.3 |
|  | 1.2 | 1.27 | 175.1 | 20.2 | 1.08 | 43.81 | 26.9 |
|  | 2 | 1.23 | 65.3 | 21.6 | 1.15 | 46.27 | 24.4 |
| 3 | 1.1 | 1.10 | 184.4 | 27.1 | 0.93 | 65.92 | 33.1 |
|  | 1.2 | 1.06 | 195.9 | 18.2 | 0.90 | 43.94 | 24.4 |
|  | 2 | 1.17 | 129.7 | 25.8 | 0.95 | 46.63 | 33.0 |
| 4 | 1.1 | 1.10 | 92.5 | 21.9 | 1.06 | 72.77 | 23.3 |
|  | 1.2 | 1.14 | 237.8 | 21.4 | 1.00 | 80.34 | 26.3 |
|  | 2 | 1.11 | 205.0 | 13.7 | 1.02 | 72.35 | 16.9 |
| 5 | 1.1 | 1.04 | 397.2 | 26.2 | 1.02 | 378.82 | 26.7 |
|  | 1.2 | 1.07 | 522.3 | 10.9 | 1.03 | 379.81 | 12.6 |
|  | 2 | - | - | - | - | - | - |
| 6 | 1.1 | 1.21 | 177.4 | 15.1 | 1.08 | 48.08 | 19.8 |
|  | 1.2 | 1.20 | 154.3 | 18.6 | 1.05 | 47.10 | 24.1 |
|  | 2 | 1.09 | 84.9 | 18.0 | 1.00 | 45.99 | 21.4 |
| 7 | 1.1 | 1.14 | 47.5 | 24.3 | 1.06 | 38.21 | 26.8 |
|  | 1.2 | 1.18 | 183.4 | 18.7 | 1.08 | 77.10 | 22.6 |
|  | 2 | 1.20 | 268.4 | 22.9 | 1.05 | 70.43 | 28.4 |
| 8 | 1.1 | 1.25 | 123.4 | 19.8 | 1.05 | 38.01 | 26.8 |
|  | 1.2 | 1.32 | 329.2 | 16.7 | 1.17 | 63.66 | 21.9 |
|  | 2 | 1.13 | 209.0 | 17.5 | 1.02 | 64.02 | 21.9 |
| 9 | 1.1 | 1.10 | 128.7 | 14.6 | 0.99 | 42.27 | 19.1 |
|  | 1.2 | 1.14 | 205.3 | 14.7 | 1.04 | 55.82 | 18.8 |
|  | 2 | 1.13 | 153.3 | 15.3 | 1.02 | 48.32 | 19.6 |
| 10 | 1.1 | 1.11 | 185.0 | 14.8 | 1.00 | 50.10 | 19.2 |
|  | 1.2 | 1.10 | 68.2 | 17.9 | 1.02 | 44.19 | 20.8 |
|  | 2 | 1.11 | 122.7 | 16.4 | 1.03 | 58.25 | 19.5 |
| 11 | 1.1 | 1.09 | 141.6 | 22.5 | 1.03 | 108.41 | 24.5 |
|  | 1.2 | 1.12 | 171.9 | 21.3 | 1.01 | 79.55 | 25.4 |
|  | 2 | 1.14 | 206.8 | 21.0 | 1.03 | 113.32 | 24.9 |
| 12 | 1.1 | - | - | - | - | - | - |
|  | 1.2 | 1.04 | 519.1 | 14.5 | 0.98 | 324.22 | 16.8 |
|  | 2 | 1.08 | 184.3 | 9.7 | 1.05 | 72.49 | 11.3 |
| 13 | 1.1 | 1.05 | 251.2 | 13.4 | 0.92 | 43.46 | 18.4 |
|  | 1.2 | 1.00 | 68.8 | 14.6 | 0.98 | 62.77 | 15.1 |
|  | 2 | 1.15 | 182.5 | 14.6 | 0.98 | 33.49 | 21.3 |
| 14 | 1.1 | 1.18 | 212.9 | 17.7 | 1.09 | 87.25 | 21.2 |
|  | 1.2 | 1.17 | 42.7 | 18.7 | 1.14 | 38.64 | 19.6 |
|  | 2 | 1.17 | 193.1 | 12.7 | 1.03 | 39.58 | 17.8 |
| 15 | 1.1 | 1.12 | 84.7 | 18.2 | 0.99 | 40.01 | 22.9 |
|  | 1.2 | 1.18 | 159.2 | 14.2 | 1.10 | 64.01 | 17.0 |
|  | 2 | 1.11 | 209.5 | 14.9 | 1.01 | 65.55 | 18.6 |
| 16 | 1.1 | 1.08 | 594.4 | 14.0 | 0.97 | 67.75 | 18.3 |
|  | 1.2 | 1.08 | 446.1 | 15.4 | 0.97 | 230.84 | 19.7 |
|  | 2 | 1.03 | 220.6 | 17.0 | 0.95 | 116.36 | 20.1 |
| 17 | 1.1 | 1.17 | 603.9 | 11.3 | 1.04 | 43.60 | 16.1 |
|  | 1.2 | 1.15 | 321.5 | 11.5 | 1.01 | 34.27 | 17.2 |
|  | 2 | - | - | - | - | - | - |
| 18 | 1.1 | 1.05 | 64.5 | 17.4 | 0.96 | 40.25 | 20.5 |
|  | 1.2 | 1.14 | 516.9 | 14.4 | 1.08 | 339.30 | 16.5 |
|  | 2 | - | - | - | - | - | - |

**Supplementary Table 2.**
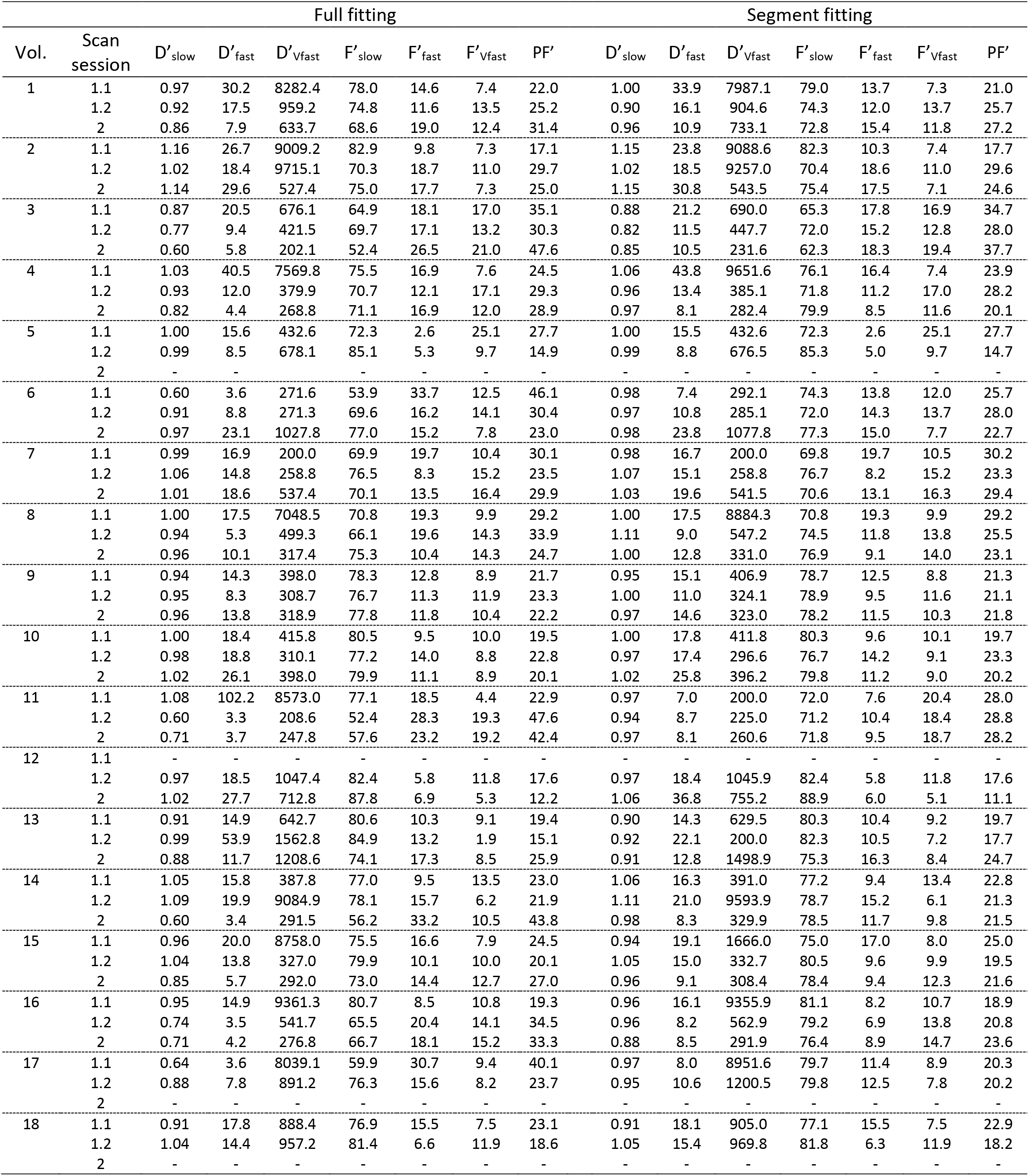
IVIM parameters estimated with tri-exponential model (Vol.: Volunteers; D’_slow_, D’_fast_, D’_Vfast_ in 10^-3^ mm^2^/s; F’_slow_, F’_fast_, F’_Vfast_, PF: %; F’_Vfast_ calculated by (1-F’_slow_ –F’_fast)_; PF’ calculated by (1- F’_slow_)).

**Supplementary Table 3.** Comparison of tri- and bi-exponential models and fitting methods using extra-sum-of-square tests (F-tests) and Akaike information criteria with the individual scans’ data (n=50).

| Models and method | Mean<br>ΔRSS | Mean<br>ΔAIC | Mean<br>ΔAICc | ΔAICc<br>(over 50 scans) | | | | | F-tests<br>p<0.05<br>(over 50 scans) |
| --- | --- | --- | --- | --- | --- | --- | --- | --- | --- |
|  |  |  |  | > 0 | 0 – 2 | 2 – 4 | 4 – 10 | >10 |  |
| Tri seg<sup>a</sup> vs. Tri full<sup>b</sup> * | 0.67 | 0.77 | 0.77 | 48 | 41* | 3 | 3* | 1* |  |
| Bi full<sup>a</sup> vs. Tri full<sup>b</sup> | 29.62 | 13.74 | 9.74 | 42 | 5 | 4 | 12 | 21 | 44 |
| Bi seg<sup>a</sup> vs. Tri full<sup>b</sup> # | 66.39 | 22.89 | 18.89 | 45 | 1 | 1 | 4<sup>#</sup> | 39<sup>#</sup> | 48 |
| Bi full<sup>a</sup> vs. Tri seg<sup>b</sup> | 28.95 | 12.97 | 8.97 | 42 | 6 | 3 | 17 | 16 | 44 |
| Bi seg<sup>a</sup> vs. Tri seg<sup>b</sup> | 65.72 | 22.12 | 18.12 | 46 | 1 | 1 | 5 | 39 | 48 |
| Bi seg<sup>a</sup> vs. Bi full<sup>b</sup> | 36.77 | 9.15 | 9.15 | 50 | 9 | 6 | 12 | 23 |  |

Tri full: Tri-exponential full fitting, Tri seg: Tri-exponential segmented fitting, Bi full: Bi-exponential full fitting, Bi full: Bi-exponential segmented fitting, AIC: Akaike Information criterion, AICc: second order Akaike Information Criterion. ΔAIC, ΔAICc difference between AIC and AICc, respectively. ΔRSS refers to the difference between residual sum-of-squares, i.e. *approach^a–^ approach^b^*. That ΔRSS is very small means there is not much difference between the two approaches. When ΔRSS is a positive value means *approach^b^* is a priori favored (with less error). When ΔAICc > 0, the *approach^b^* is more likely to be correct than the *approach^a^;* with weak evidence and considerable evidence when 0 ≤ ΔAICc ≤ 2 and 4 ≤ ΔAICc ≤ 7, respectively. With ΔAICc ≥ 10, the *approach^a^* is considered unlikely to be correct compared to *approach^b^* (17). *; as an example, according to ΔAICc, there are 41 scans slightly favor Tri full, while only 4 scans strongly favor Tri full (ΔAICc > 4, as compared with Tri-seg). #; as an example, according to ΔAICc, there are 43 scans strongly favor Tri-full (ΔAICc > 4, as compared with Bi-seg). F-tests p<0.05: numbers of scans where *extra-sum-of-squares F-tests* were significant between nested models with p<0.05, that is, the test result shows the more complex approach, i.e. *tri-exponential* is justified (15).

**Supplementary Table 4.** Comparison of tri- and bi-exponential models and fitting methods using adjusted R^2^ and information criteria with total-averaged 50 scans liver signal.

| Models | Adjusted-R<sup>2</sup><br>(×10<sup>-2</sup>) | RSS | AIC | AICc | ΔAICc<sub>1</sub> | ΔAICc<sub>2</sub> | Evidence Ratio |  |
| --- | --- | --- | --- | --- | --- | --- | --- | --- |
|  |  |  |  |  |  |  | Tri full vs.<br>other<br>methods | Bi full vs.<br>Bi seg |
| Bi-exponential |  |  |  |  |  |  |  |  |
| Seg | 98.57 | 70.11 | 77.05 | 80.68 | 65.81 | 14.21 | 1.95×10<sup>14</sup> | 1.22×10<sup>3</sup> |
| Full | 99.41 | 28.84 | 62.83 | 66.47 | 51.60 | 0.00 | 1.60×10<sup>11</sup> | - |
| Tri-exponential |  |  |  |  |  |  |  |  |
| Seg | 99.98 | 0.81 | 5.69 | 15.02 | 0.15 | - | 1.08 | - |
| Full <sup>#</sup> | 99.98 | 0.80 | 5.54 | 14.87 | 0.00 | - | - | - |

Seg: Segmented fitting, Full: full fitting, Tri full: tri-exponential full fitting, Bi full: bi-exponential full fitting, Bi seg: bi-exponential segmented fitting. RSS: Residual sum of squares, AIC: Akaike Information Criterion, AICc: Second order Akaike Information Criterion (RSS, AIC, AICc: the smaller the better). ΔAICc_**1**_ means the difference in AICc between a model and the model with the lowest AICc of the set (^#^ tri-exponential full fitting presented the lowest AICc and is therefore considered as reference). ΔAICc_2_ means the difference in AICc between bi-exponential segmented fitting and full fitting model. Evidence Ratio (Tri full vs. other methods) corresponds to the relative likelihood between the best model (^#^ tri-exponential full fitting) and other models, meaning how many times tri-exponential full fitting model is more likely to be correct than the other models. Evidence Ratio (Bi full vs. Bi seg) corresponds to the relative likelihood between bi-exponential full and segmented fitting methods. RSS analysis shows tri-exponential model is a priori favored over bi-exponential model, and for bi-exponential model full fitting is favored over segmented fitting. ΔAICc_**1**_ and evidence ratio show bi-exponential model is unlikely to be correct compared to tri-exponential full fitting. However, for tri-exponential model, ΔAICc between full fitting and segmented fitting is very small. Full fitting is only 1.08 times more likely to be correct than segmented fitting according to evidence ratio. For bi-exponential model, ΔAICc_2_ shows that segmented fitting is unlikely to be correct compared with full fitting. According to evidence ratio full fitting is more than ≥1000 times more likely to be correct than segmented fitting.

**Supplementary Figure 1.**
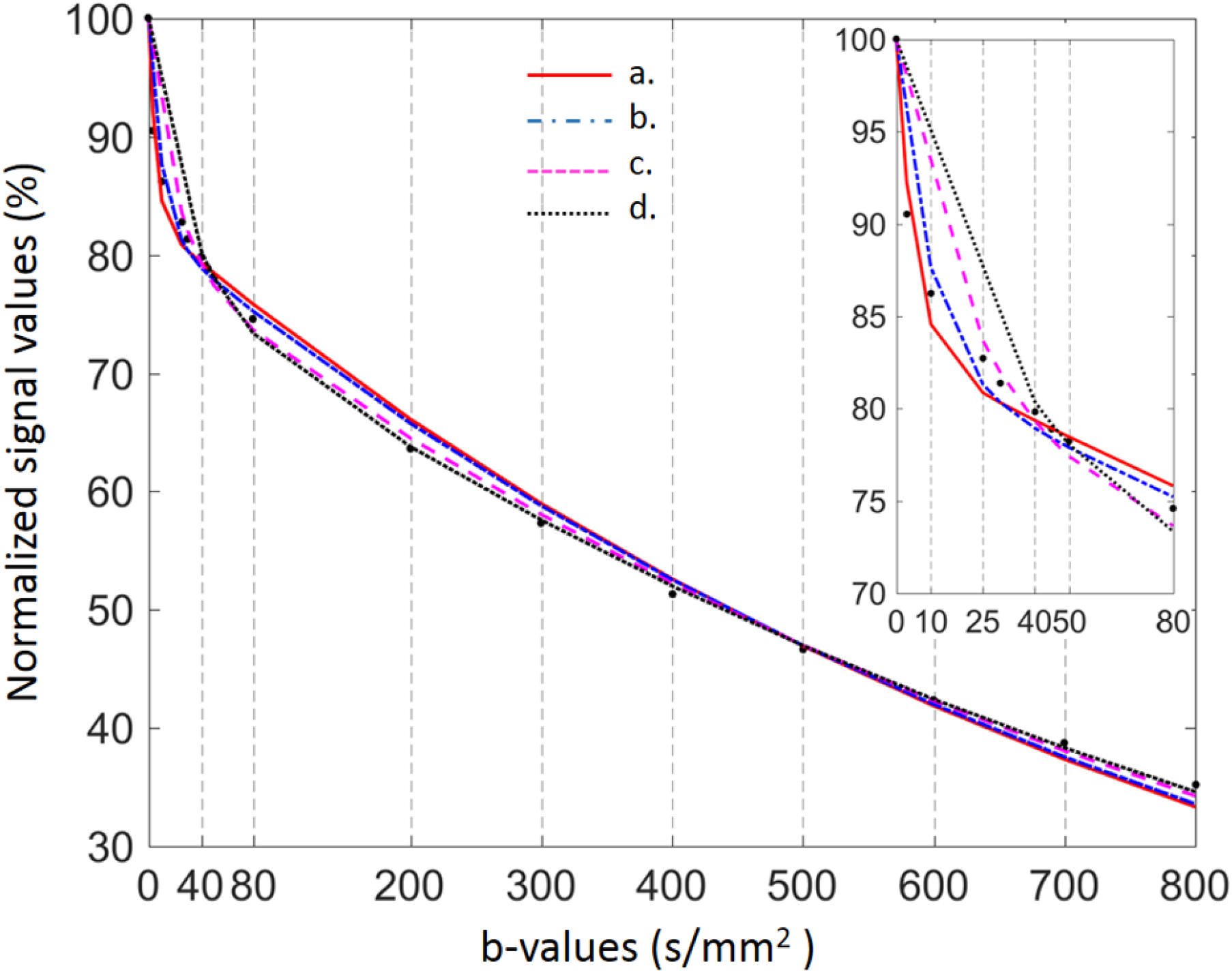
A comparison of total-averaged signal fitting with bi-exponential full fitting using 16 *b*-values distribution (0, 3, 10, 25, 30, 40, 45, 50, 80, 200, 300, 400, 500, 600, 700, 800 s/mm^2^)(***a***) and after the successive removals of very low and low *b*-values (***b**, **c**, **d***). After the successive removals of very low and low *b*-values, the initial slope of the fitted-curve becomes less and less steep, D_fast_ value therefore decreases. D_slow_ value also decreases, while PF value increases. (***b***: *b*=3 s/mm^2^ removed; ***c***: *b*=3, 10 s/mm^2^ removed, ***d***: *b*=3, 10, 25, 30 s/mm^2^ removed. D_fast_=193.6, 103.6, 48.1, and 35.2 ×10^-3^ mm^2^/s, D_slow_=1.14, 1.12, 1.06, and 1.02 ×10^-3^ mm^2^/s, PF=16.9%, 17.7%, 20.3%, and 21.8%, for ***a, b, c, d***, respectively)

**Supplementary Figure 2.**
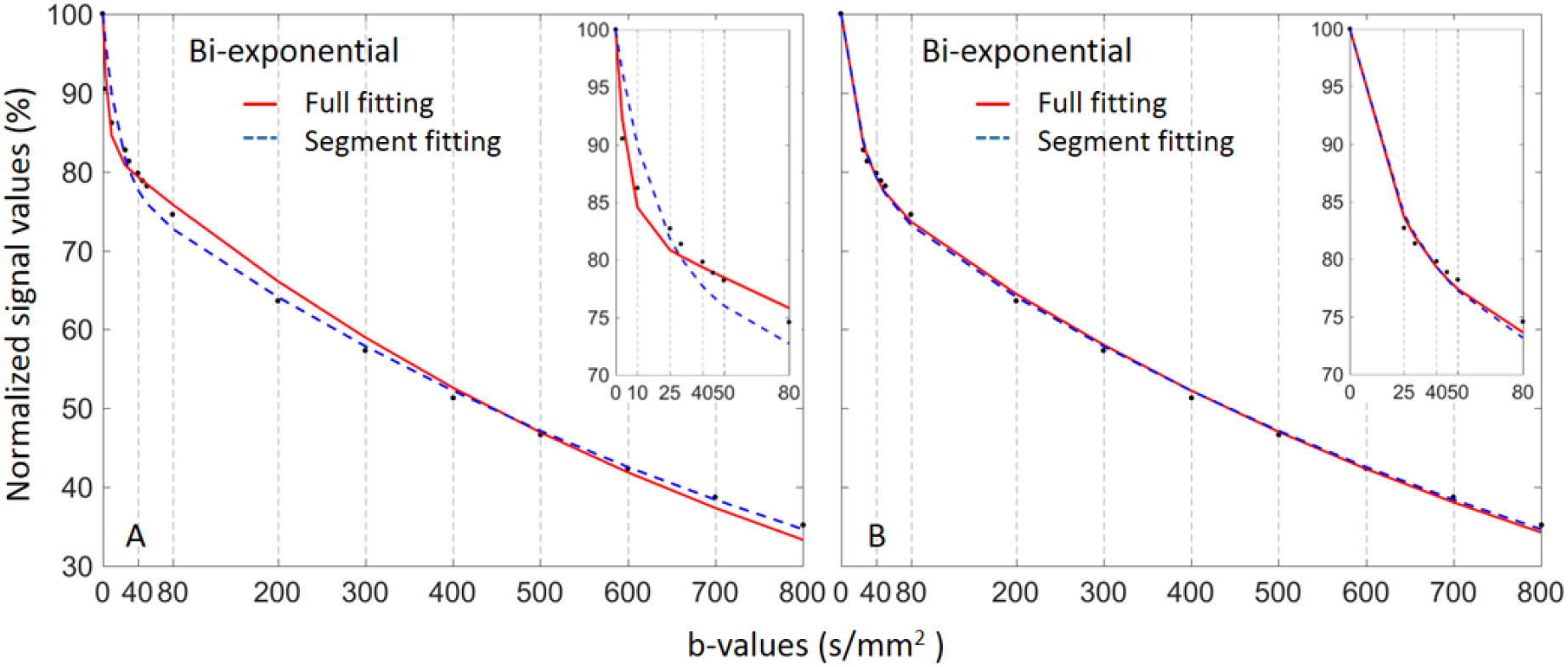
A comparison of total-averaged signal fitting using bi-exponential model with full or segmented fitting, using 16 *b*-values distribution (A) and after the removal of very low *b*-values (*b*=3, 10 s/mm^2^) (B). The IVIM parameters estimated from (A)&(B) are listed in the table below. The removal of very low *b*-values leads to lower D_fast_ values, especially for full fitting method, since the initial fast decay is almost totally removed. It also leads to lower D_slow_ and higher PF values for full fitting. For segmented fitting, both PF and D_slow_ remain the same, since D_slow_ and PF are calculated first (signal value at *b*=0 remains unchanged). Therefore, the distribution of very low *b*-value has more impact with full fitting than with segmented fitting. Note that in the current example the fitted-curves and IVIM parameters derived from full and segmented fittings are more similar after the removal of very low *b*-values.

| Fitting method | $b$-values distribution | $D_{\text{fast}} \times 10^{-3}$ mm<sup>2</sup>/s | $D_{\text{slow}} \times 10^{-3}$ mm<sup>2</sup>/s | PF in % |
| --- | --- | --- | --- | --- |
| full fitting | full 16 $b$-values (A) | 193.6 | 1.14 | 16.9 |
| | limited low $b$-values (B) | 48.1 | 1.06 | 20.3 |
| Seg fitting | full 16 $b$-values (A) | 56.7 | 1.03 | 21.3 |
| | limited low $b$-values (B) | 42.9 | 1.03 | 21.3 |

## References

1. Le Bihan D, Breton E, Lallemand D, et al. Separation of diffusion and perfusion in intravoxel incoherent motion MR imaging. Radiology. 1988;168:497–505.

2. Zhang Q, Wang YX, Ma HT, et al. Cramér-Rao bound for Intravoxel Incoherent Motion Diffusion Weighted Imaging fitting. Conf Proc IEEE Eng Med Biol Soc 2013;2013:511–514.

3. Li YT, Cercueil JP, Yuan J, et al. Liver intravoxel incoherent motion (IVIM) magnetic resonance imaging: a comprehensive review of published data on normal values and applications for fibrosis and tumor evaluation. Quant Imaging Med Surg 2017;7:59–78.

4. Chevallier O, Zhou N, He J, et al. Removal of evidential motion contaminated and poorly fitted image data improves IVIM diffusion MRI parameter scan–rescan reproducibility. Acta Radiologica DOI: 10.1177/0284185118756949

5. Park HJ, Sung YS, Lee SS, et al. Intravoxel incoherent motion diffusion-weighted MRI of the abdomen: The effect of fitting algorithms on the accuracy and reliability of the parameters. J Magn Reson Imaging 2017;45:1637–1647.

6. Cercueil JP, Petit JM, Nougaret S, et al. Intravoxel incoherent motion diffusion-weighted imaging in the liver: comparison of mono-, bi- and tri-exponential modelling at 3.0-T. Eur Radiol 2015;25:1541–50.

7. Wurnig MC, Germann M, Boss A. Is there evidence for more than two diffusion components in abdominal organs? – A magnetic resonance imaging study in healthy volunteers. NMR in Biomedicine. 2018;31:e3852. https://doi.org/10.1002/nbm.3852

8. Turner R, Le Bihan D, Maier J, et al. Echo-planar imaging of intravoxel incoherent motion. Radiology 1990;177:407–414.

9. Li Y, Lu P-X, Huang H, et al. Dependence of Intravoxel Incoherent Motion MR threshold b-value selection for separating perfusion and diffusion compartment and liver fibrosis diagnostic performance. bioRxiv [Internet]. 2017 Jul 15; Available from: http://biorxiv.org/content/early/2017/07/15/164129.abstract

10. Barbieri S, Donati OF, Froehlich JM, et al. Impact of the calculation algorithm on biexponential fitting of diffusion-weighted MRI in upper abdominal organs. Magn Reson Med 2016;75:2175–2184.

11. Dillon WR, Goldstein M. Multivariate Analysis–Methods and Applications. New York: Wiley, 1984

12. Li YT, Huang H, Zhuo Z, et al. Bi-phase age-related brain gray matter magnetic resonance T1ρ relaxation time change in adults. Magn Reson Imaging 2017;39:200–205.

13. Wang YXJ, Deng M, Li YT, et al. A Combined Use of Intravoxel Incoherent Motion MRI Parameters Can Differentiate Early-Stage Hepatitis-b Fibrotic Livers from Healthy Livers. SLAS Technol 2017 doi: 10.1177/2472630317717049

14. Lemke A, Laun FB, Simon D, et al. An in vivo verification of the intravoxel incoherent motion effect in diffusion-weighted imaging of the abdomen. Magn Reson Med. 2010 Dec;64(6):1580–5.

15. Motulsky H, Christopoulos A. Fitting Models to Biological Data Using Linear and Nonlinear Regression: A Practical Guide to Curve Fitting. London: Oxford Univ. Press, 2004

16. Yuan J, Wong OL, Lo GG, et al. Statistical assessment of bi-exponential diffusion weighted imaging signal characteristics induced by intravoxel incoherent motion in malignant breast tumors. Quant Imaging Med Surg. 2016;6:418–429.

17. Burnham KP, Anderson DR. Multimodel Inference: Understanding AIC and BIC in Model Selection. Sociol Methods Res. 2004;33:261–304.

18. Gurney-Champion OJ, Froeling M, Klaassen R, et al. Minimizing the Acquisition Time for Intravoxel Incoherent Motion Magnetic Resonance Imaging Acquisitions in the Liver and Pancreas. Invest Radiol 2016;51:211–20.

19. Luciani A, Vignaud A, Cavet M, et al. Liver cirrhosis: intravoxel incoherent motion MR imaging--pilot study. Radiology 2008;249:891–899.

20. Andreou A, Koh DM, Collins DJ, et al. Measurement reproducibility of perfusion fraction and pseudodiffusion coefficient derived by intravoxel incoherent motion diffusion-weighted MR imaging in normal liver and metastases. Eur Radiol 2013;23:428–434.

21. Cohen AD, Schieke MC, Hohenwalter MD, et al. The effect of low b-values on the intravoxel incoherent motion derived pseudodiffusion parameter in liver. Magn Reson Med 2015;73:306–311.

22. Hayashi T, Miyati T, Takahashi J, et al. Diffusion analysis with triexponential function in liver cirrhosis. J Magn Reson Imaging 2013;38:148–53.

23. Hayashi T, Miyati T, Takahashi J, et al. Diffusion analysis with triexponential function in hepatic steatosis. Radiol Phys Technol. 2014;7:89–94.

24. Pazahr S, Nanz D, Rossi C, et al. Magnetic resonance imaging of the liver: apparent diffusion coefficients from multiexponential analysis of b values greater than 50 s/mm2 do not respond to caloric intake despite increased portal-venous blood flow. Invest Radiol. 2014;49:138–46.

## Suppl Doc References

1. Jerome NP, d’Arcy JA, Feiweier T, et al. Extended T2-IVIM model for correction of TE dependence of pseudo-diffusion volume fraction in clinical diffusion-weighted magnetic resonance imaging. Phys Med Biol 2016;61:N667–N680.

2. Dyvorne H, Jajamovich G, Kakite S, et al. Intravoxel incoherent motion diffusion imaging of the liver: optimal b-value subsampling and impact on parameter precision and reproducibility. Eur J Radiol 2014 83:2109–2113.

3. Park HJ, Sung YS, Lee SS, et al. Intravoxel incoherent motion diffusion-weighted MRI of the abdomen: The effect of fitting algorithms on the accuracy and reliability of the parameters. J Magn Reson Imaging 2017;45:1637–1647.

4. Barbieri S, Donati OF, Froehlich JM, et al. Impact of the calculation algorithm on biexponential fitting of diffusion-weighted MRI in upper abdominal organs. Magn Reson Med 2016;75:2175–2184.

5. Cohen AD, Schieke MC, Hohenwalter MD, et al. The effect of low b-values on the intravoxel incoherent motion derived pseudodiffusion parameter in liver. Magn Reson Med 2015;73:306–311.

6. Luciani A, Vignaud A, Cavet M, et al. Liver cirrhosis: intravoxel incoherent motion MR imaging--pilot study. Radiology 2008;249:891–899.

7. Dyvorne H, Jajamovich G, Kakite S, et al. Intravoxel incoherent motion diffusion imaging of the liver: optimal b-value subsampling and impact on parameter precision and reproducibility. Eur J Radiol 2014 83:2109–2113.

8. Patel J, Sigmund EE, Rusinek H, et al. Diagnosis of cirrhosis with intravoxel incoherent motion diffusion MRI and dynamic contrast-enhanced MRI alone and in combination: preliminary experience. J Magn Reson Imaging 2010;31:589–31600.

9. Guiu B, Petit J-M, Capitan V, Aho S, et al. Intravoxel Incoherent Motion Diffusion-weighted Imaging in Nonalcoholic Fatty Liver Disease: A 3.0-T MR Study. Radiology. 2012;265:96–103.

10. Regini F, Colagrande S, Mazzoni LN, et al. Assessment of Liver Perfusion by IntraVoxel Incoherent Motion (IVIM) Magnetic Resonance-Diffusion-Weighted Imaging: Correlation With Phase-Contrast Portal Venous Flow Measurements. J Comput Assist Tomogr. 2015;39:365–72.

11. Hu F, Yang R, Huang Z, et al. Liver fibrosis: in vivo evaluation using intravoxel incoherent motion-derived histogram metrics with histopathologic findings at 3.0 T. Abdom Radiol (NY). 2017;42:2855–63.

12. Luo M, Zhang L, Jiang X-H, Zhang W-D. Intravoxel Incoherent Motion Diffusion-weighted Imaging: Evaluation of the Differentiation of Solid Hepatic Lesions. Transl Oncol. 2017;10:831–8.

13. Wurnig MC, Donati OF, Ulbrich E, et al. Systematic analysis of the intravoxel incoherent motion threshold separating perfusion and diffusion effects: Proposal of a standardized algorithm. Magn Reson Med. 2015;74:1414–22.

14. Li Y, Lu P-X, Huang H, Leung J, Chen W, Wang Y-X. Dependence of Intravoxel Incoherent Motion MR threshold b-value selection for separating perfusion and diffusion compartment and liver fibrosis diagnostic performance. bioRxiv [Internet]. 2017 Jul 15; Available from: http://biorxiv.org/content/early/2017/07/15/164129.abstract

15. Chevallier O, Zhou N, He J, Loffroy R, Wáng YX. Removal of evidential motion contaminated and poorly fitted image data improves IVIM diffusion MRI parameter scan–rescan reproducibility. Acta Radiologica DOI: 10.1177/0284185118756949

16. Cercueil JP, Petit JM, Nougaret S, et al. Intravoxel incoherent motion diffusion-weighted imaging in the liver: comparison of mono-, bi- and tri-exponential modelling at 3.0-T. Eur Radiol 2015;25:1541–50.

17. Li YT, Cercueil JP, Yuan J, et al. Liver intravoxel incoherent motion (IVIM) magnetic resonance imaging: a comprehensive review of published data on normal values and applications for fibrosis and tumor evaluation. Quant Imaging Med Surg 2017;7:59–78.

18. Dillon WR, Goldstein M. Multivariate Analysis–Methods and Applications. New York: Wiley, 1984

19. Yin P, Fan X. Estimating R 2 Shrinkage in Multiple Regression: A Comparison of Different Analytical Methods. The Journal of Experimental Education. 2001;69:203–24.

20. Motulsky H, Christopoulos A. Fitting Models to Biological Data Using Linear and Nonlinear Regression: A Practical Guide to Curve Fitting. London: Oxford Univ. Press, 2004

21. Burnham KP, Anderson DR. Multimodel Inference: Understanding AIC and BIC in Model Selection. Sociol Methods Res. 2004;33:261–304.

